# A Bundled Antiparallel Cytochrome Nanowire Structure Suggests Roles in Cell-Cell Electron Transfer and Biofilm Formation

**DOI:** 10.1101/2025.10.22.684050

**Authors:** Holly A. Petersen, Chi H. Chan, Gilbert O. Carpenter, Media Zakizadeh Tabari, Jessie Lynda Fields, Ayisha Zia, Shane T. Rich-New, Allon I. Hochbaum, Daniel R. Bond, Fengbin Wang

## Abstract

Long-range extracellular electron transfer enables respiring microbes to use minerals, other organisms, or electrodes as electron acceptors by transporting electrons microns away from the cell surface. This process is primarily studied in *Geobacter sulfurreducens*, which produces at least three different micrometer-long, multi-heme cytochrome nanowires capable of electron transfer. However, the distribution and higher-order structure of these types of cytochrome nanowires remains largely unknown. Here, we employed cryo-electron microscopy to determine the atomic structure of a unique cytochrome nanowire from *Desulfuromonas soudanensis* WTL, a halophilic, iron- and electrode-reducing bacterium found in deep subsurface brine. These filaments are based on a homolog of the OmcE tetraheme cytochrome, which then assemble into highly ordered bundles of antiparallel filaments. This arrangement likely arises from the association of nanowires extending from adjacent cells. Furthermore, a similar cytochrome bundle structure was observed in *Geobacter metallireducens*, suggesting that this quaternary structure may be a common feature among nanowires secreted by electroactive microbes. Our findings demonstrate that cytochrome nanowires in diverse taxa can form specialized bundle interfaces, potentially facilitating conductive biofilm formation and representing a novel strategy for microbial electron exchange. More broadly, this work expands understanding of electron transfer mechanisms and demonstrates the production of multi-heme filaments across distinct lineages. These insights could guide future research into cytochrome nanowire secretion and conductive biofilm engineering, with potential applications in electrochemical technologies.

## INTRODUCTION

In environments lacking oxygen, many respiring prokaryotes rely on extracellular electron transfer to move electrons beyond their outer membrane to a terminal electron acceptor^1,2^. First observed in the reduction of iron and manganese oxides^3,4^, extracellular electron transfer also allows cells to dispose of reducing equivalents using nearby syntrophic partners such as methanogens^5,6^. These organisms still generate ATP through electron transport across the cytoplasmic membrane, but must direct electrons from the quinone pool to periplasmic carriers and ultimately across the insulating outer membrane to reach extracellular chelated minerals, oxide particles, organisms, and surfaces^7–9^. Some organisms rely primarily on diffusive redox shuttles such as flavins to carry electrons from outer membrane multi-heme cytochromes, as observed in *Shewanella*^10–12^. However, if soluble metals or electron shuttles are not readily available, redox proteins must physically contact the acceptor surfaces, which can be microns away^13–15^. To accomplish this, organisms such as *Geobacter sulfurreducens*^16–21^ produce extracellular helical conductive appendages.

*G. sulfurreducens* has been used as a model system for studying bacterial extracellular electron transfer^22–24^. Initial analysis of the genome showed it encodes a remarkable number of multiheme cytochrome genes, including 70+ open reading frames with three or more CxxCH heme c binding motifs, and for decades these cytochromes were hypothesized to contribute to electron transfer only in the inner membrane, periplasm, and outer membrane. Cryogenic electron microscopy (cryo-EM) revealed that at least three different multiheme *c*-type cytochromes in *G. sulfurreducens* polymerize into micrometer-long appendages: OmcS^19,21^, OmcE^17^, and OmcZ^18,20^. Although these three proteins contain only about 20% secondary structure and share little structural or sequence similarity with each other, they can be classified into two categories based on the arrangement of hemes within the assembled cytochrome filaments: “insulated” and “leaky” nanowires^20^. OmcS and OmcE have their heme molecules well-insulated by surrounding protein backbones. This type of insulated cytochrome nanowire, with identical heme arrangements but completely different subunit structure, has also been observed in Archaea^25^, such as *Pyrobaculum calidifontis* and *Archaeoglobus veneficus*.

In contrast, OmcZ has a significantly different heme arrangement. Each subunit contains a branched, solvent-exposed “leaky” heme in addition to the linear conductive chain of hemes. Without this exposed heme, the linear chain would be insulated, resembling OmcS and OmcE. As the *omcZ* gene is upregulated 400–800% in electrode-associated biofilms^26,27^, and its deletion impairs anodic electron transfer to electrodes^27–29^, networks of OmcZ filaments are proposed to form an electron grid to make a conductive biofilm, providing multiple pathways for electron flow^18^. Cryo-EM visualization of sheared OmcZ did reveal many nanowires clumped together, but without specific, repeating interactions^18,20^. To date, no atomic structures showing direct interactions between any class of nanowire has been reported.

Microbial appendages are common components of biofilms, along with other biomaterials such as exopolysaccharides and extracellular DNA^30^. In fact, appendages often drive biofilm formation^31–33^. For example, retractable bacterial type IV pili are critical for adherence to surfaces, ultimately leading to colonization and biofilm formation^34^. Appendage-driven biofilm formation can also involve formation of specific pilus-to-pilus interfaces, resulting in the formation of ordered pili bundles. One such bundle structure was resolved by cryo-EM in *Pyrobaculum calidifontis*^35^, where the pili assemble into highly ordered bipolar bundles. The bipolar nature of these bundles likely arises from the association of pili from at least two different cells. Pili important in biofilm formation have also been observed to form bundles in bacteria, such as TasA^36^ in *Bacillus subtilis* and the chaperone-usher pathway (CUP) pilus CupE^37^ in *Pseudomonas aeruginosa*. However, only single filament structures were resolved in these cases; therefore, the pilus-pilus interfaces within the bundles, whether specific or not, remain unresolved.

In this work, we present the cryo-EM structure of a bundle formed by cytochrome nanowires produced by *Desulfuromonas soudanensis* WTL, a halophilic iron-reducing bacterium isolated from anoxic deep subsurface brine. This is the first structure of a cytochrome nanowire in a Bacterium outside of *G. sulfurreducens*. We show that the primary cytochrome building block is homologous to *G. sulfurreducens* OmcE. Our structural data clearly reveal that *D. soudanensis* OmcE nanowires form antiparallel bundles mediated by two types of filament-to-filament interactions. Unlike *P. calidifontis*, which produces two different appendages, including one non-bundle-forming cytochrome filament and another bundling pilus for biofilm formation, *D. soudanensis* WTL produces cytochrome bundles that may be capable of both electron transfer and cell-cell association functions. In-depth sequence analysis identified likely OmcE homologs in five different bacterial phyla and revealed highly variable sequence regions within the OmcE subunit. We demonstrate that this bundling phenomenon is not unique, showing that OmcE nanowires also form bundles in *Geobacter metallireducens*, a bacterium evolutionarily much closer to *G. sulfurreducens*. Such large, ordered bundles of nanowires are of great interest in protein engineering, offering potential biosensing applications such as detection of viruses and tumor biomarkers^38–40^. Furthermore, by combining this with structural analysis of heme arrays from over 500 AlphaFold^41^ predicted OmcE-like proteins, we provide insights into understanding these dual-function cytochrome nanowire bundles.

## RESULTS

### *D. soudanensis* WTL produces bundles of OmcE cytochrome nanowires

In order to investigate *D. soudanensis* WTL extracellular appendages, we grew 1 L of *D. soudanensis* WTL cells^42^ and enriched extracellular filaments by ammonium sulfate precipitation as previously described^43^. Cryo-EM micrographs of the enriched extracellular filaments showed that they formed filament bundles with diameters up to 150 nm, while single filaments were sparsely observed (Figure 1A). To validate that the bundle formation was physiologically relevant, and not introduced by the enrichment buffer, we examined sheared filaments under two other conditions: phosphate-buffered saline, and the bacterial growth medium^42^ (containing a total concentration of over 500 mM of various salts that mimic the natural brine, including CaCl2, MgCl2, and NaCl). Strikingly, large numbers of filamentous bundles were observed under all tested conditions, and single filaments were rarely seen (Supplemental Figure 1), suggesting that the sheared appendages interact and bundle together regardless of salinity.

**Figure 1.**
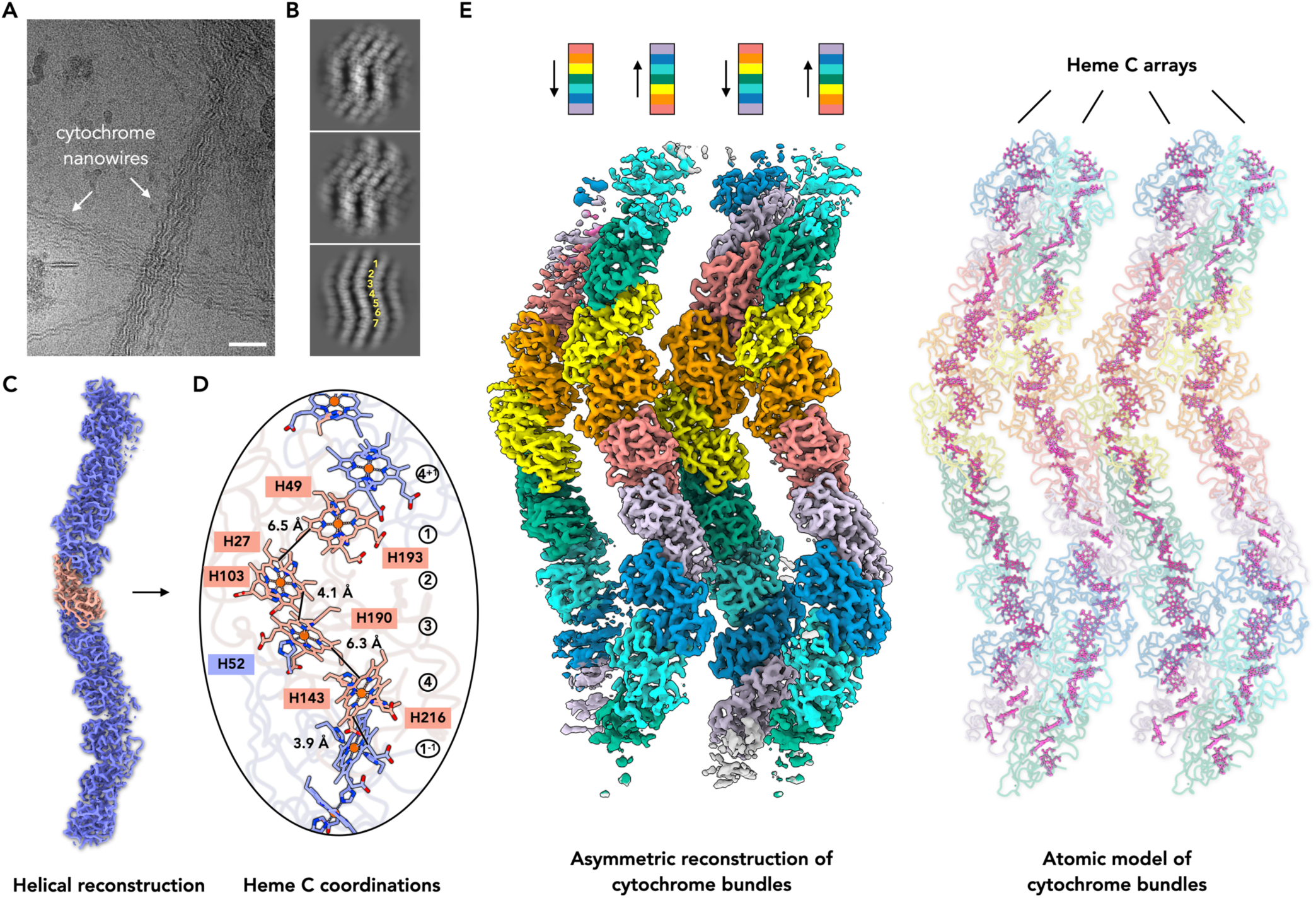
*D. soudanensis* WTL produces bundles of OmcE cytochrome nanowires. (A) Representative cryo-EM micrograph of cytochrome bundles from D. soudanensis WTL. Scale bar: 50 nm. (B) 2D class averages of cytochrome bundles. (C) Helical reconstruction of a single cytochrome filament within the cytochrome bundles, with one cytochrome subunit highlighted in salmon. (D) Arrangement of heme molecules within the helical reconstruction. (E) Asymmetric reconstruction of cytochrome bundles and the corresponding atomic model built into the map. To clearly demonstrate the filament’s polarity, seven different subunit conformations are colored using a rainbow scheme, as indicated. The protein backbone of the atomic model is colored to match the map, while all heme molecules are colored magenta.

After data collection, the resulting 2D averages revealed only seven distinct views along the filament (Figure 1B), suggesting a preferred orientation problem, a rare occurrence in helical reconstructions. Indexing the power spectrum of the bundles indicated a helical twist of ∼51.4 degrees and a rise of ∼30.8 Å (Supplemental Figure 2), confirming the observations in the 2D classes. To obtain additional views of the filaments, tilted data were collected with the sample grid tilted at -12°, +12°, and +4°, respectively. With the inclusion of these tilted micrographs, helical reconstruction of the single cytochrome filament reached a resolution of 3.1 Å, as determined by map-to-map FSC (Supplemental Figure 2).

The cytochrome building block was identified as DSOUD_2954, a homolog of the tetraheme OmcE^17^ first reported in *G. sulfurreducens*. Consistent with the structures of previously reported “insulated” cytochrome nanowires in *G. sulfurreducens*^17,19^ and two Archaeal species^25^, the orientation between adjacent hemes was either stacked or T-shaped, with minimum distances between porphyrins ranging from 3.9 to 6.5 Å (Figure 1D). On the surface of the WTL OmcE, two ion-binding sites were observed. Based on the coordinating distances^44^ and medium components, these ions are likely to be calcium, which possibly stabilizes the cytochrome nanowires under high salt conditions (Supplemental Figure 3).

After helical averaging, only a single cytochrome fiber could be seen in the cryo-EM volume, suggesting that the bundle lacked global helical symmetry. We then asked the question of how individual cytochrome filaments interact to form these large bundles. To avoid introducing reference bias, we used the aligned particles and volumes obtained after helical refinement to perform a 3D classification with 10 classes. If multiple specific interfaces existed between single fibers within the bundle, these different classes could then be sorted out and reconstructed independently.

Following 3D classification, several classes revealed new, ordered cytochrome filaments adjacent to the starting single filament reference. Strikingly, manual inspection of all these volumes consistently showed the same anti-parallel packing arrangement between cytochrome nanowires (Figure 1E). This difference in polarity between adjacent filaments within the same bundle suggests they likely originate from different bacterial cells, and the inter-filament interfaces are specific. These 3D classes were subsequently merged for an asymmetrical reconstruction, which reached a 3.1 Å resolution as judged by map-to-map FSC (Supplemental Figure 2).

This 3D classification revealed the presence of new, ordered cytochrome filaments adjacent to the starting single filament reference. Strikingly, manual inspection of these volumes consistently showed the same anti-parallel packing arrangement between cytochrome nanowires (Figure 1E). This alternating polarity between adjacent filaments within the same bundle suggested the inter-filament interfaces were specific. These 3D classes were subsequently merged for an asymmetrical reconstruction, which reached a 3.1 Å resolution as judged by map-to-map FSC (Supplemental Figure 2).

### *D. soudanensis* bundles are anti-parallel arrays of tightly packed cytochrome nanowires

Since the asymmetrical reconstruction resolved a large area within the bundle and the helical parameters of single filaments, we could expand the atomic model to better visualize the entire filament array (Figure 2A). As indicated by the helical twist, 2D classification, and 3D reconstruction, the cytochrome subunit circles back to an identical environment every seven subunits (Figure 1E, 2A). Furthermore, the 3D reconstruction revealed that every other cytochrome filament in the bundling array not only has the same polarity but is also in register (Figure 2A). Therefore, to examine inter-filament interfaces, we only needed to consider the relationships within these seven distinct cytochrome subunit states, using a single cytochrome filament as a reference. These seven states are visualized by drawing imaginary lines perpendicular to the bundle direction (Figure 2A). Intriguingly, interfaces were observed for only four of these seven states (Figure 2B); in states where filaments were not sufficiently close to form detectable contacts, the amino acid configuration was different (Figure 2C).

**Figure 2.**
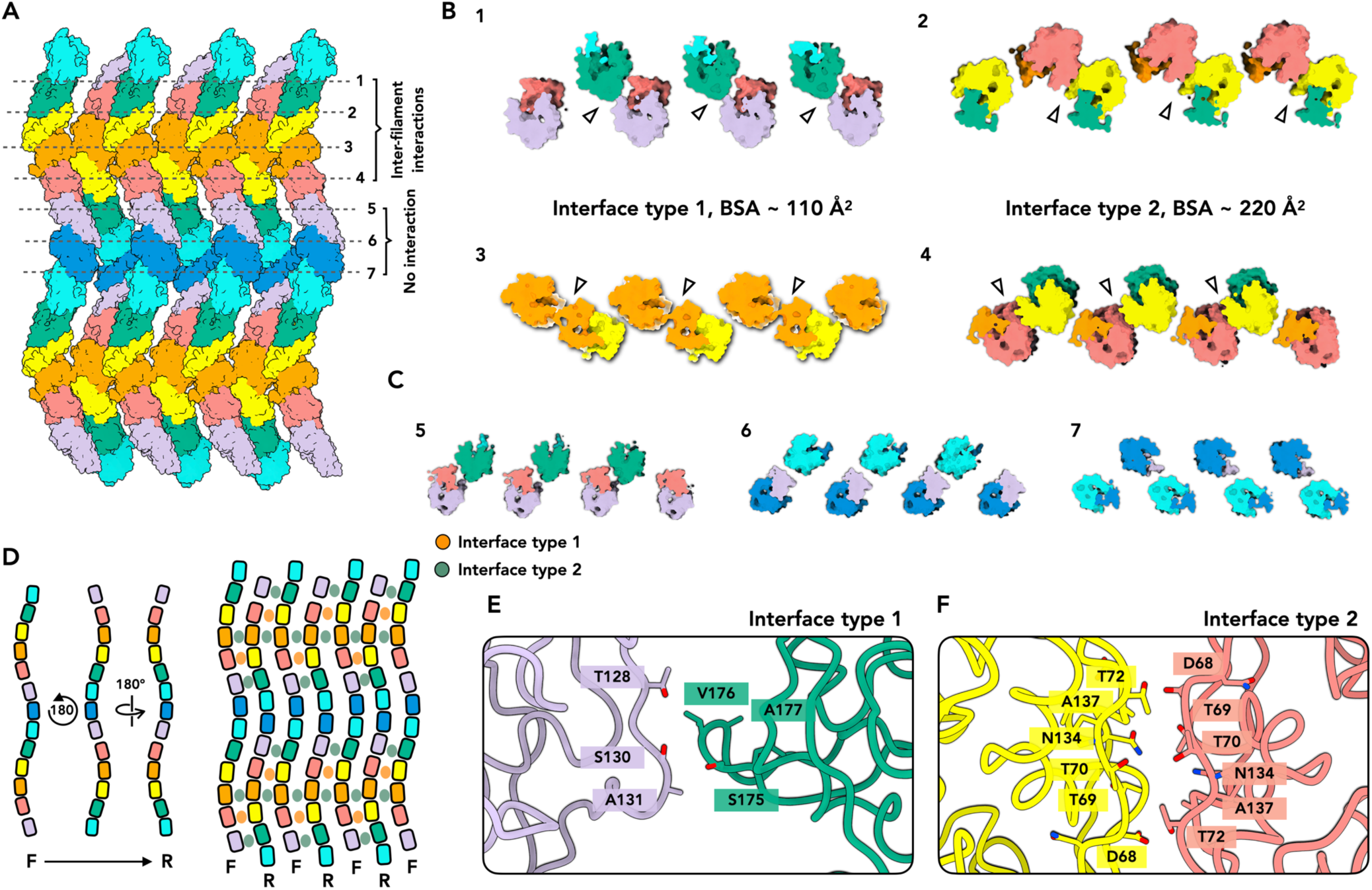
Binding interface within anti-parallel arrays of cytochrome nanowires. (A) Expanded atomic model of cytochrome arrays based on the obtained cryo-EM map. Different subunit conformations are colored using the same rainbow scheme as in Figure 1. Seven unique inter-filament conformations are indicated by dashed lines. (**B-C**) The seven unique inter-filament conformations corresponding to those indicated in (A) are viewed from the top. Two types of interfaces are shown in (B), while three conformations without evident inter-filament interfaces are shown in (C). (**D**) A schematic illustrating the anti-parallel arrangement of cytochrome bundles and the locations where the interfaces are observed. (**E-F**) Zoomed-in views of the type 1 and type 2 interfaces.

Where inter-filament contacts were detected, two unique interfaces were observed, with buried surface areas of 110 Å^2^ (type 1 interface) and 220 Å^2^ (type 2 interface), as estimated by PISA^45^. Despite being small, these interfaces accumulate rapidly along the bundles (Figure 2D). For instance, between two adjacent cytochrome filaments, the interface area is 440 Å^2^ per 21.7 nm. This scales to a substantial interface of approximately 101,000 Å^2^ for two 5 µm long cytochrome filaments, and this is amplified further in a bundling array containing dozens of such long filaments. When looking at the interfaces more closely, the type 1 interface is asymmetric, involving two loop regions: T128-A131 from one subunit and S175-A177 from another subunit (Figure 2E). In contrast, the type 2 interface is symmetric, primarily mediated by the same loop regions, D68-T72 and N134-A137, between two subunits (Figure 2F).

Given the presence of seven distinct states of cytochrome subunits along a single cytochrome fiber, we then investigated their dynamic behavior. Notably, when applying a global helical symmetry during reconstruction, several loop regions appeared smeared and less defined in the resulting map. However, upon relaxing the helical symmetry in the asymmetrical reconstruction, these loops became clearly traceable (Figure 3A-B), suggesting that they are ordered loop structures that adopt different conformations across the seven states. This observation was further validated by calculating the RMSD between the seven subunits, which revealed that these same loop regions undergo moderate conformational changes (Figure 3C). Intriguingly, the loops exhibiting conformational changes overlap with the type 1 interface (T128-A131) and are in close proximity to a portion of the type 2 interface (N134-A137) (Figure 3D).

**Figure 3.**
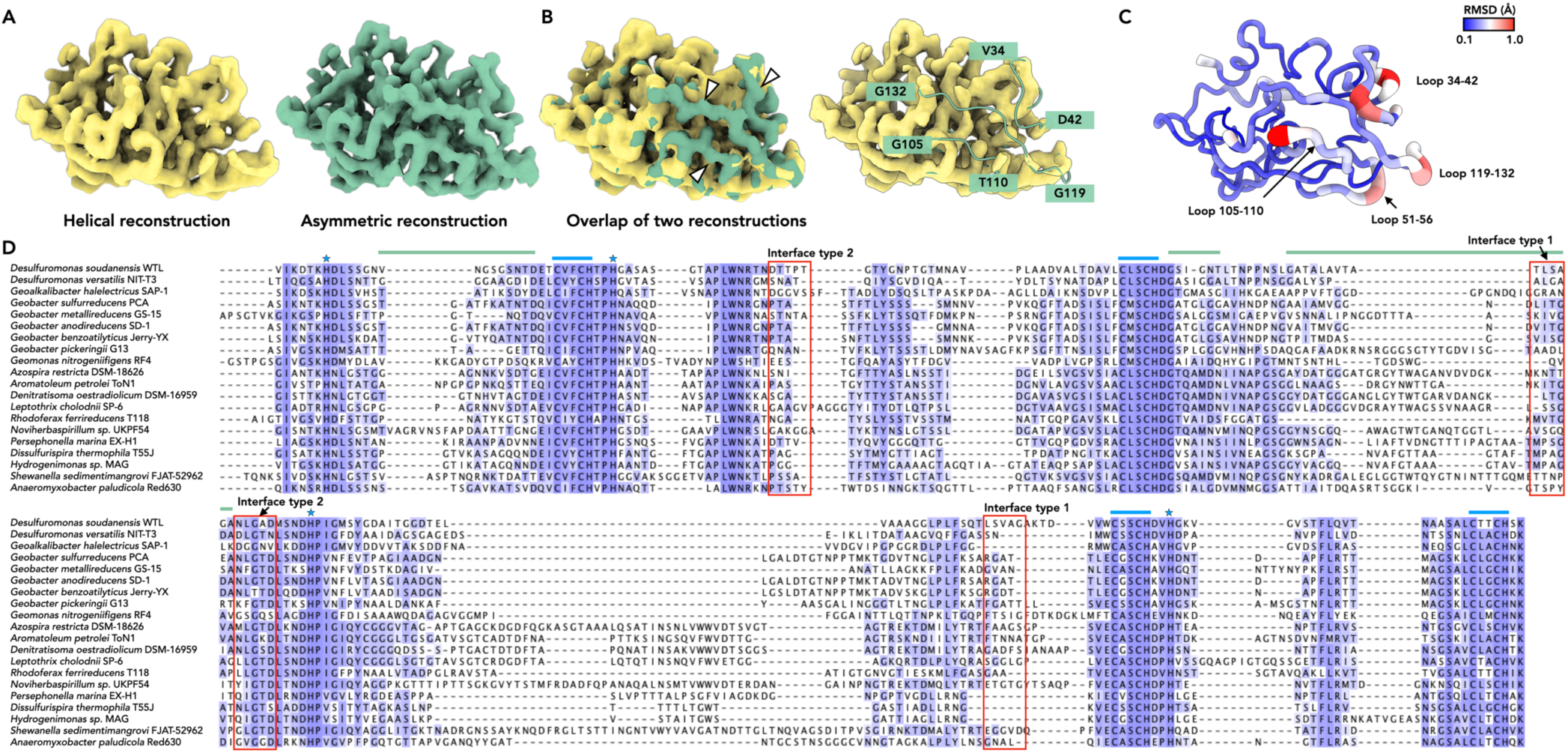
Loops involved in WTL bundling interface are not conserved among OmcE homologs. (**A**) Cryo-EM maps of a single D. soudanensis WTL OmcE monomer, obtained from helical refinement (left) and asymmetric refinement (right). (**B**) Overlay of the two volumes from (A) reveals that several loop regions in the helical refinement are smeared out when helical symmetry is applied, suggesting they have different conformations. (**C**) RMSD analysis of seven different cytochrome conformations from asymmetric refinement indicates that similar loop regions have adopted slightly different conformations. (**D**) Structure-based sequence alignment of OmcE homologs from various bacteria. All selected OmcE ORFs have an outer membrane porin gene located nearby. Conserved heme-binding motifs and histidines are indicated by blue lines and stars, respectively. Loop regions involved in the *D. soudanensis* WTL bundling interface are highlighted by red boxes. Loop regions that adopt different conformations within D. soudanensis WTL bundles are indicated by green lines.

### *D. soudanensis* bundle interface loops are not conserved among OmcE homologs

As the bundling of cytochrome nanowires is a newly observed phenomenon, we were curious whether the amino acids participating in the specific bundling interfaces, or within the loops exhibiting conformational changes, were conserved among other OmcE homologs. Considering the widespread distribution of tetraheme cytochromes with sequences similar to OmcE, we narrowed our sequence analysis to OmcE-like proteins that are potentially capable of assembling into nanowires by only including OmcE homologs where other genes characteristic of this nanowire occurred, such as gene encoding a large outer membrane ß-barrel protein putatively functioning as a secretion or anchor, and a ß-propeller domain lipoprotein (see Figure 4).

**Figure 4.**
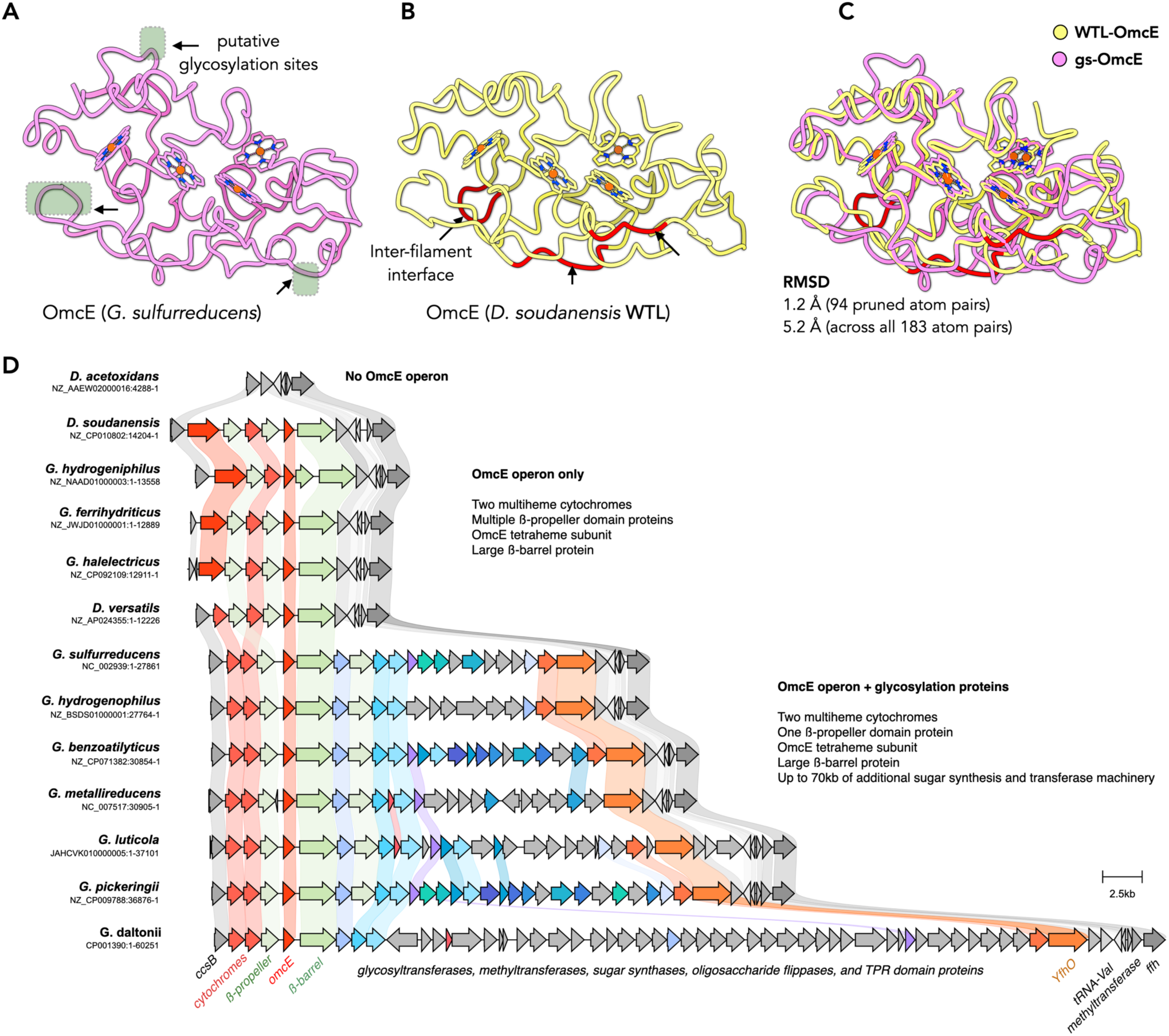
D. soudanensis WTL OmcE vs. G. sulfurreducens OmcE. (**A**) A single OmcE subunit from the cryo-EM structure of the *G. sulfurreducens* OmcE filament. Regions where additional cryo-EM densities were observed are indicated by green squares. (**B**) A single OmcE subunit from the bundled structures of *D. soudanensis* WTL OmcE. Loop regions involved in inter-filament interfaces are colored in red. (**C**) Superimposition of the two structures shown in (A) and (B). (**D**) Comparison of the *omcE* gene cluster between *G. sulfurreducens*, *D. soudanensis* WTL, and other bacterial strains.

Remarkably, putative OmcE homologs were identified across five different phyla, predominantly in Thermodesulfobacteriota and Pseudomonadota, with some examples also found in Aquificota, Campylobacterota, and Nitrospirota. A sequence alignment of homologs from these five phyla revealed several conserved regions, including the four CxxCH motif regions and the other four histidine residues involved in heme coordination. However, strikingly, almost none of the *D. soudanensis* loops that facilitate in bundle formation were conserved among any subset of these OmcE homologs (Figure 3D). The only region displaying moderate conservation is N134-A137, which is part of the type 2 interface; however, this limited conservation might be attributed to its relatively close proximity to a conserved histidine residue that coordinates internal heme.

### Similarities between OmcE nanowire structures

As the *D. soudanensis* nanowire bundle is the second experimentally characterized OmcE nanowire, we investigated its differences from the *G. sulfurreducens* version. The amino acid sequence of *D. soudanensis* OmcE exhibits 41% sequence identity to *G. sulfurreducens* OmcE, and both structures are primarily composed of random coil domains. Similar to *G. sulfurreducens*, *D. soudanensis* OmcE contains ∼3% β-strands and ∼17% α-helices, with the remaining 80% adopting coil and turn conformations. While the overall architectures of these two cytochromes appear similar, conformational variations are noticeable (Figure 4A-B). In *G. sulfurreducens* OmcE, extra densities, likely corresponding to glycosylation, were observed^17^. In contrast, the cryo-EM map of *D. soudanensis* OmcE did not reveal such densities, suggesting an absence of post-translational modifications.

Consistent with this observation, the loops containing putative glycosylation sites in *G. sulfurreducens* OmcE are considerably different from those in WTL OmcE. Furthermore, the loops implicated in inter-filament interfaces in the OmcE bundle structure also exhibit distinct conformations compared to their counterparts in *G. sulfurreducens* (Figure 4A-C). This divergence is reflected in the structural alignment of the two cytochromes (Figure 4C). The RMSD between 94 pruned atom pairs, encompassing all heme-coordinating regions, is ∼1.2 Å, indicating a conserved heme packing core. However, upon expanding the alignment to include all ∼180 atom pairs, the RMSD increases to 5.2 Å, highlighting significant conformational differences in non-heme coordinating regions. These differences are largely attributable to the distinct surface residues, namely those involved in glycosylation versus inter-filament interaction.

As glycosylation appeared to be a major difference between these structures, we compared genomic regions across the Desulfuromonadia (Figure 4D). Nearly all genomes possess a region of shared synteny beginning with purine metabolism (*purHDE)* and heme biosynthesis (*resB*, *ccsB)*, and ending with tRNA(Val)-modification and signal recognition particle (*ffh)*. Some genomes encode an OmcE-containing cluster between these genes. The smallest clusters always encode two mutiheme cytochromes, two ß-propeller lipoproteins, the OmcE nanowire subunit, and a large ß-barrel protein. This pattern is primarily found in marine, alkaliphilic, and thermophilic genera (classically named *Desulfuromonas, Geoalkalibacter, and Geothermobacter).* A second and more common pattern contains the same OmcE gene cluster, followed by 15 to 70 genes encoding glycosyltransferases (typically families 2, 4 and 39), sugar modification enzymes, methyltransferases, sugar exporters, tetratricopeptide domain proteins, and transcriptional regulators. These additional glycosylation genes are found primarily in genera associated with freshwater and subsurface habitats, such as *Geobacter* and *Geomonas.* These genomic differences are consistent with our observed lack of glycosylation in *D. soudanensis*.

### Heme arrangements in OmcE-like structures and a possible new OmcE nanowire family

All known ‘insulated’ cytochrome nanowires, including OmcE^17^ and OmcS^19^ from *G. sulfurreducens*, pcECN from *P. calidifontis*, and avECN^25^ from *A. veneficus*, share a similar arrangement of densely packed hemes. Within this arrangement, the porphyrin rings of heme-heme pairs are in either “parallel” or “T-shaped” alternating configurations. Despite their significant structural and sequence differences, the heme arrangement in the *D. soudanensis* OmcE filament is nearly identical to that of *G. sulfurreducens*.

Furthermore, AlphaFold predictions of all other OmcE homologs suggest that they also share the same heme arrangement (Figure 5A). A meta-analysis of heme pairs in all multiheme c-type cytochrome structures deposited in the Protein Data Bank reveals that the vast majority of these pairs are arranged in parallel or T-shaped configurations^25^. When we added *D. soudanensis* OmcE to this analysis (Figure 5B), it clearly fell within these two preferred regions.

**Figure 5.**
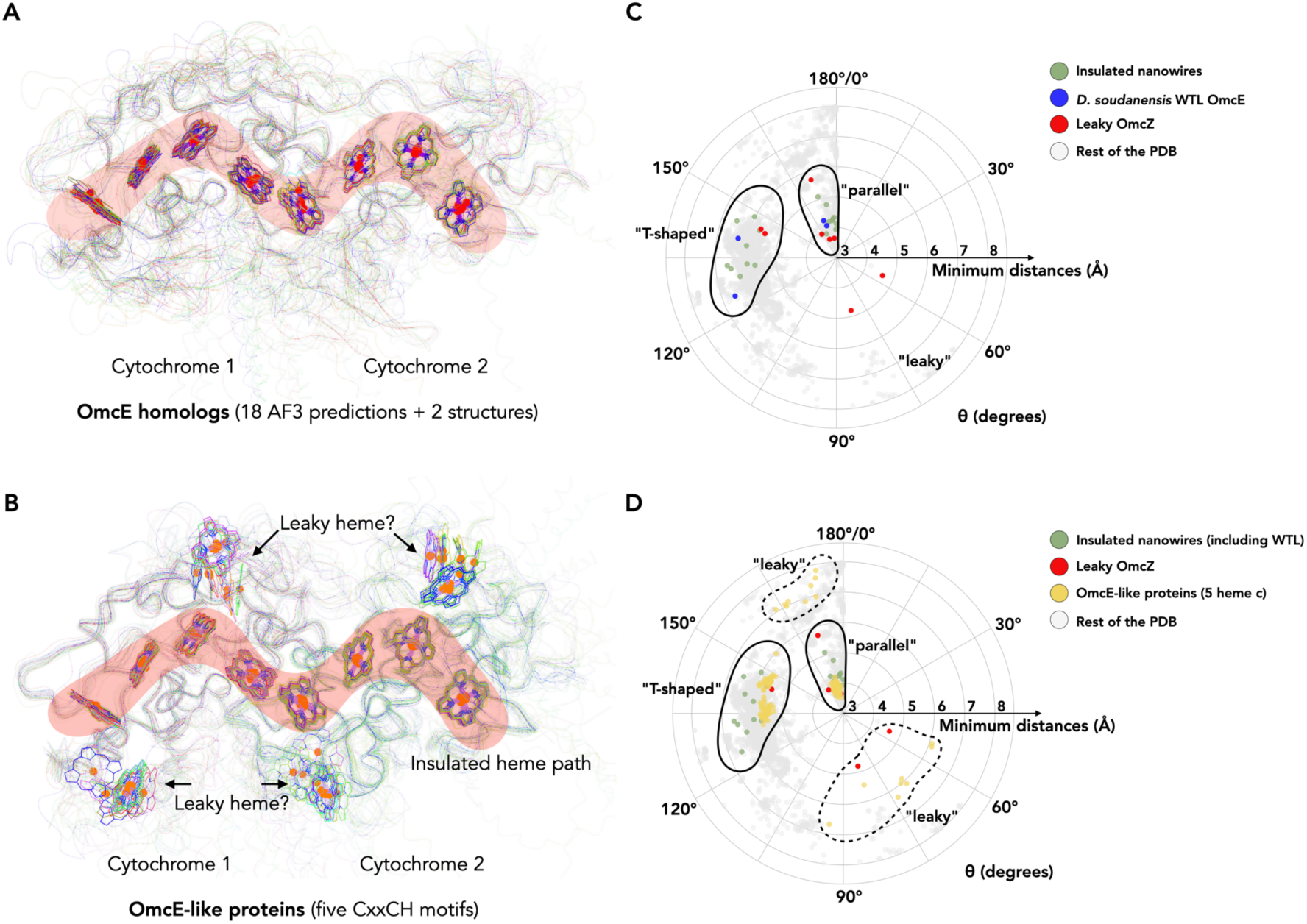
Heme arrangements in OmcE-like proteins. (**A**) Two subunits of OmcE homologs (sequence listed in Figure 3). Experimentally determined OmcE models were used for *D. soudanensis* WTL and *G. sulfurreducens*. Other OmcE proteins were predicted using AlphaFold3 with heme molecules and aligned to the *D. soudanensis* WTL OmcE model. Heme molecules are shown as sticks, while all protein backbones are displayed with very thin ribbons. (**B**) Similar to (A), but instead of OmcE homologs, this panel shows protein predictions with sequence similarity to OmcE but possessing five CxxCH motifs. These predictions were aligned to the experimentally determined *D. soudanensis* WTL OmcE model. (**C**) Heme-heme orientation plot. One heme can be aligned to an adjacent heme by a rotation and a translation. Only the 25 non-hydrogen atoms in the porphyrin ring were used in the alignment. Heme pairs in all PDB structures were analyzed. Minimum distances refer to the shortest distance between two porphyrin rings, regardless of atom type. The angle θ was determined from the alignment rotation matrix between heme pairs. All porphyrin ring pairs with a minimum distance less than or equal to 8 Å are shown. Porphyrin ring pairs in insulated nanowires, the *D. soudanensis* WTL OmcE, and the “leaky” OmcZ are shown in green, blue, and red, respectively. (**D**) Heme-heme orientation plot similar to (C). Porphyrin ring pairs in insulated nanowires (including *D. soudanensis* WTL OmcE), “leaky” OmcZ, and OmcE-like proteins from (B) are shown in green, red, and yellow, respectively.

The broadly conserved geometry of these heme pairs suggests a convergent design principle has evolved to allow electron transfer through multiheme c-type cytochromes. It is worth noting that all measured rotation angles for the “parallel” heme pair configuration are between 170 and 180 degrees, because the vinyl groups of these heme pairs are positioned on opposite sides of their porphyrin ring planes, resulting in a rotation angle close to 180 degrees, rather than 0 degrees, when the two heme molecules are aligned.

Interestingly, when we searched for OmcE-like proteins via PSI-BLAST, we identified many proteins with additional CxxCH binding motifs, despite maintaining a similar protein length. Most of these proteins belong to uncultured bacterial species distributed across diverse phyla. We were curious about the potential location of these extra hemes, especially if these proteins were to assemble into a nanowire. Therefore, we aligned AlphaFold predictions of these proteins to subunits of the *D. soudanensis* OmcE model (Figure 5C). Remarkably, these 5-heme OmcE-like proteins still exhibit the conserved insulated heme arrangement, while the additional heme is much more solvent-accessible. The extra heme also appears to have a preferable packing location (Figure 5C). This observation is reminiscent of the “leaky” nanowire OmcZ^18^ from *G. sulfurreducens*, where each cytochrome subunit has a solvent-accessible heme. We then plotted the heme-heme pairs of these OmcE-like proteins on the polar plot. Indeed, we observed two additional clusters originating from the extra heme: one cluster closely resembling that of known OmcZ, and another cluster similar to the “stacked” pairs but with greater porphyrin-porphyrin distances (Figure 5D). This suggested the existence of an uncharacterized nanowire family with properties similar to both OmcE and OmcZ.

### Bundling of OmcE cytochrome nanowires: a common phenomenon?

The bundling effects observed in *D. soudanensis* OmcE are not unique. We also observed bundle formation in extracellular appendages sheared from *Geobacter metallireducens* (Supplemental Figure 4), an iron-reducing bacterium^46^ often studied as a partner in direct interspecies electron transfer^47^. *G. metallireducens* is a freshwater isolate phylogenetically closer to *G. sulfurreducens* than to *D. soudanensis*. Notably, *G. metallireducens* does not encode OmcS, but it does encode OmcE and OmcZ. Further, the gene cluster in *G. metallireducens* is highly similar to *G. sulfurreducens,* and encodes a clear glycosylation cluster.

Using cryo-EM to analyze the *G. metallireducens* extracellular appendages, we identified type IV pili made of PilA-N and PilA-C and putative cytochrome nanowires. These cytochrome nanowires were present both as single filaments and in large bundles. 2D class averages of the cytochrome bundles showed specific interfaces between the filaments, allowing resolving multiple filaments in one average (Supplemental Figure 4). The average power spectrum revealed a helical symmetry for a single cytochrome nanowire of 29 Å rise and 60° twist, consistent with known symmetries for OmcE nanowires, as the rise and twist are much larger for OmcZ and OmcS filaments. Furthermore, the protein subunits within these 2D classes displayed the “Pac-Man” shape consistent with other experimentally determined OmcE structures (Supplemental Figure 4). While the limited number of orientations resulted in only six views, preventing reconstruction of the bundle to near-atomic resolution, our observations confirm bundling of OmcE nanowires occurs in *G. metallireducens* as well. If this quaternary arrangement is more common than previously recognized, future cryo-EM studies need to take this into consideration, and examine bundles as well as individual filaments.

## DISCUSSION

Recent advances in cryo-EM^48^ and deep-learning methods^49,50^ have significantly reshaped the field of microbial electron transfer, and current techniques have proven very valuable for *de novo* identification of extracellular protein polymers without prior knowledge^51–53^. In this study, we used cryo-EM to examine the sheared total appendages from the iron-reducing bacterium *D. soudanensis* WTL^42^. Strikingly, the only type of filaments observed were thick bundles formed by anti-parallel packed multi-heme c-type cytochromes, based on a subunit homologous to *G. sulfurreducens* OmcE.

This is the first instance of a Bacterial cytochrome nanowire structure being experimentally determined outside of *G. sulfurreducens*. Furthermore, we showed that another organism, *G. metallireducens,* also produces cytochrome filaments in bundles consistent with the same OmcE subunit, indicating such bundling is not a unique case. The unexpected observation of bundling in cytochrome nanowires suggests possible roles in colonization and biofilm formation, and reveals cytochrome nanowire structures can be more complex than initially anticipated. It also provides insights into how such architecture might facilitate electron transfer between cells.

As previously reported^42^, *D. soudanensis* WTL grows more slowly than many better-known metal-reducing bacteria, with a doubling time of approximately 13 hours. For an organism isolated from a 10°C low-nutrient metal-rich brine, where significant energy is likely devoted simply to maintenance and ion homeostasis^54,55^, the large quantity of extracellular cytochrome bundles represents a substantial energy expenditure. Heme-rich *c*-type cytochromes are energetically expensive to synthesize^56^, as a filament could contain over 10,000 protein subunits and 40,000 heme cofactors, and each bacterium appears to produce at minimum several dozen such nanowires. Why *D. soudanensis* is investing a large amount of energy into this single part of its respiratory chain is an open question.

How cytochrome nanowires are secreted and assembled on the cell surface also remains unanswered. Even among bundling pili that share a similar ‘jelly-roll’ pilin fold (*P. calidifontis* archaeal bundling pili, *B. subtilis* TasA, and *P. aeruginosa* CupE), the secretion systems likely differ. The secretion mechanisms for the first two remain unclear, whereas *P. aeruginosa* CupE clearly utilizes a chaperone-usher pathway^57–59^, with the pilus mounted on an outer membrane usher β-barrel porin. In *G. sulfurreducens*, the insulated cytochrome nanowire OmcS was initially proposed to be pumped out via the ATP-dependent type IV pilus, analogous to the type II secretion system^60^. However, a more recent hypothesis suggests it is instead mounted on the outer membrane using a porin-like pathway aided by several other proteins^61^. Considering the large number of cytochrome nanowires produced by each cell, a porin model appears more energetically efficient, and consistent with the presence of large β-barrel proteins in most OmcE and OmcS genomic regions.

These questions of secretion illustrate how cryo-EM nanowire structures resolve some questions about electron transfer, while simultaneously raising many more involving physiology. For example, if OmcE assembly is via simple polymerization analogous to chaperone-usher systems, how is it terminated? In the CUP pili assembly system, the process is halted by the incorporation of a slightly different pilin. This prevents the chaperone from dissociating, thereby blocking the addition of further major pilins^62^. A similar “stopper” protein might operate in *D. soudanensis*. Because the interface between OmcE subunits within the filament features an unsatisfied heme requiring an additional histidine, an ideal stopper protein would possess nine histidines and four hemes, with a structure resembling OmcE. Using AlphaFold3, we identified one protein, Dsoud_2415, that meets these criteria in *D. soudanensis* WTL. However, this protein is not located within the same gene cluster as OmcE, while stopper proteins typically co-evolve with the major filament subunit, as observed in CUP pili^63,64^, making this hypothesis a bit less likely. Similar to the problem of termination is one of initiation– what is the structure of the filament’s tip? Could it be a protein similar to OmcE, but with more solvent-exposed hemes to enhance contact with minerals? No such candidate appears encoded in the OmcE operons.

Ultimately, what could be the advantage of a cell producing bundles rather than individual cytochrome filaments? A simple explanation is that bundling could protect filaments from degradation or phage recognition, or protect them against mechanical shearing. However, considering the anti-parallel structure, this suggests a driving force for evolution of bundling was cell-cell interaction (Figure 6). If a portion of the biofilm is in direct contact with minerals, daughter cells are more likely to remain linked simply due to the bundled nanowires keeping cells close. A more intriguing possibility involves potential electron transfer between cells. The heme-to-heme distance between the heme cores or each filament is at least 25 Å, which is too great for efficient electron transfer between nanowires in the bundles. Cell-to-cell electron transfer would be possible if the end of each nanowire makes contact with a neighboring cell. This interaction would create a highly insulated pathway for one cell to transfer electrons to a distant acceptor, using another as a transfer station (Figure 6).

**Figure 6.**
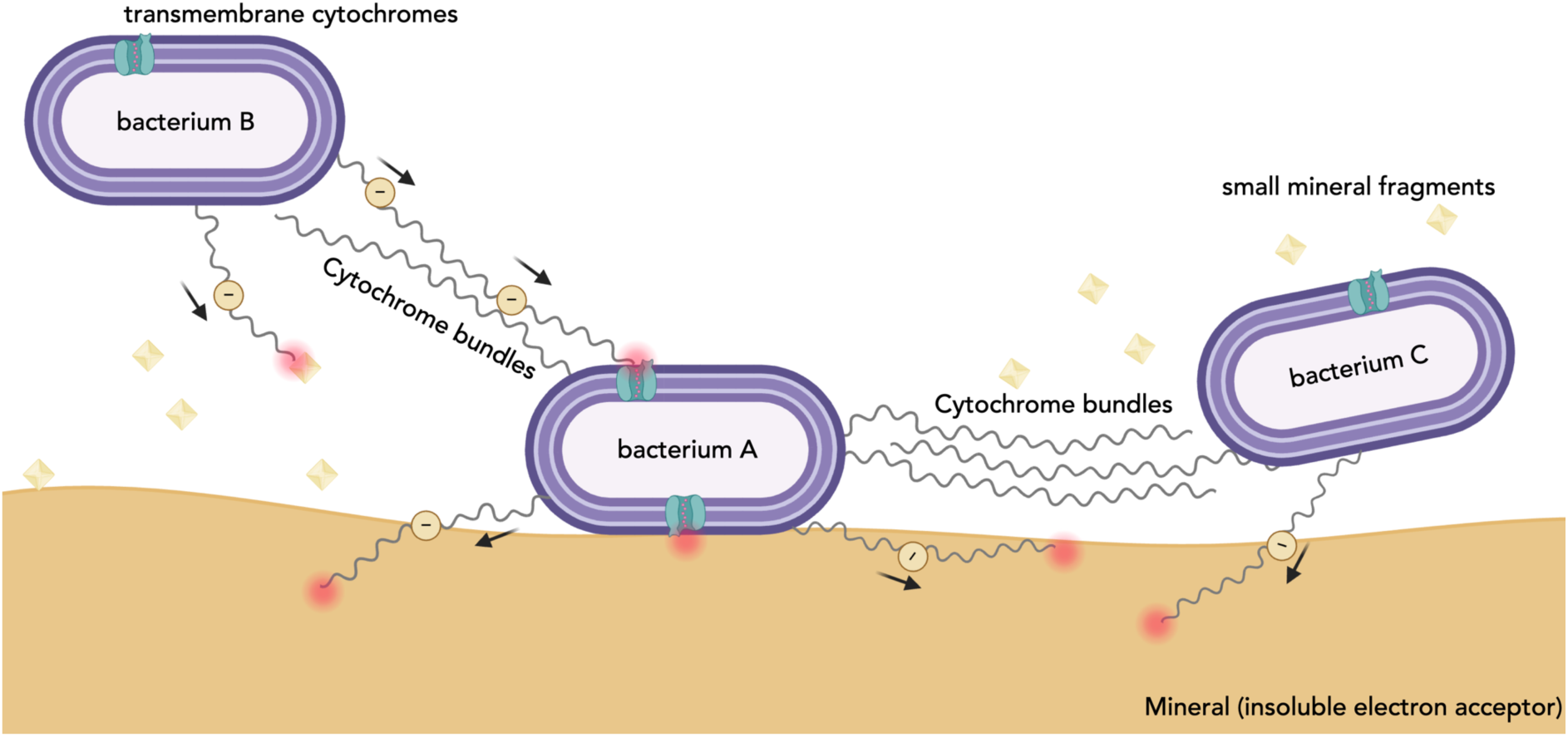
Schematic of potential cytochrome bundle functions within *D. soudanensis* WTL biofilms. Three representative bacterial cells (A-C) are depicted. OmcE nanowires are represented by thin, curved lines, and simplified bundles are indicated. Putative transmembrane or membrane-embedded cytochromes are represented by green cylinders. Minerals and mineral fragments are shown in yellow. Potential electron transfer events are indicated by a red halo effect.

In summary, our study has identified a new higher order structure for the OmcE cytochrome nanowire, based on anti-parallel bundles found in both *D. soudanensis* WTL and *G. metallireducens*. The near-atomic resolution structure of *D. soudanensis* WTL bundles suggests these nanowires could serve a dual function in both electron transfer and biofilm formation, implying a more complex role in anaerobic respiration than previously understood. This work suggests cytochrome nanowires as a widespread tool utilized in bacteria beyond *G. sulfurreducens* and highlights the need for future research to elucidate the mechanisms of cytochrome nanowire secretion, assembly, and function.

## METHODS

### *D. soudanensis* WTL cytochrome nanowire preparation

*D. soudanesis* cells were collected from 1 L of culture in a modified Soudan Mine (SM) medium. The medium was degassed in 80:20 N_2_:CO_2_ and contained 20 mM sodium acetate, 40 mM sodium fumarate, 10 ml/L mineral mixture^23^, 0.04 mM magnesium sulfate, 19 mM ammonium chloride, 0.37 mM potassium phosphate monobasic, 75 mM calcium chloride, 37 mM magnesium chloride, and 135 mM sodium chloride. The pH was adjusted to 6.8 before adding 21.5 mM sodium bicarbonate^42^. When the density reached 0.6 of OD_600_, cell culture was harvested by centrifugation for 20 minutes at 8000 x *g*. Cytochrome filament abundance and bundle formation were examined by negative staining (Supplemental Figure 5). Extracellular filaments were sheared by drawing and expelling the cell suspension through a 5 ml syringe attached to a 25g x 7/8” long needle 10 times. The debris was subsequently removed by centrifugation for 20 minutes at 8000 x *g*. The sheared filaments was then resuspended in 150 mM ethanolamine buffer, pH 10.5, before cryo-EM imaging.

### *Geobacter metallireducens* cytochrome nanowire preparation

*G. metallireducens* biofilm were grown on 9 cm^2^ graphite electrodes poised at 0 V vs. SHE were in 100 ml bioreactors in mineral medium with only 40 mM sodium acetate, without sodium nitrate, Fe citrate or sodium tungstate^23^. To increase the biomass of the biofilm, bioreactors were supplemented with 5 successive rounds of 40 mM acetate supplementation when electrode current decreased to near 0 amps. Biofilm was scraped off the graphite and resuspended in 50 mM Tris buffer at pH 7.5 with 0.25 U/ml benzonase. The filaments were sheared off by drawing and expelling cell suspension through a 5 ml syringe attached with a 25g x 7/8” long needle 10 times. The debris was removed by centrifugation for 20 minutes at 8000 x *g*.

### Negative staining electron microscopy

For negative staining imaging, continuous carbon grids (EMS, CF300-Cu) were glow-discharged for 30 s at 30 mA, with the carbon side facing up. Subsequently, 3.5 μL medium culture containing *D. soudanensis* WTL cells was directly applied to the carbon side of the glow-discharged grid and allowed to incubate for 20 s. The grid was then back-blotted by placing the copper side on filter paper (Whatman) and gently dragging downward against the filter paper to remove the excess sample. Immediately following this, 10 μL of ultrapure water was dropped onto the grid surface and quickly back-blotted as before; this washing step was repeated once. Following the second wash, 8 μL of 2% uranyl acetate (EMS) was placed onto the grid and back-blotted using the same procedure. The grid was then allowed to air dry for 1 min. TEM images were collected on the 200 keV JEOL JEM-1400 transmission electron microscope, which was equipped with an AMT-NanoSprint43L-MarkII camera, at the University of Alabama at Birmingham’s High Resolution Imaging Facility.

### Cryo-EM conditions and image processing

For cryo-EM imaging, the enriched cytochrome nanowire sample (4.5 μL) was applied to glow-discharged, lacey carbon grids and then plunge-frozen using an EM GP2 Plunge Freezer (Leica). Cryo-EM micrographs were collected on a 300 keV Titan Krios microscope equipped with a K3 camera at a pixel size of 1.07 Å and a total dose of 50 e⁻/Å². Preliminary cryo-EM analysis using a non-tilted dataset indicated only seven distinct views of cytochrome subunits within the cytochrome bundles. Therefore, we performed additional tilted data collections at +12° and -12° for grids with the thinnest ice and collected an additional dataset on a thicker ice grid using +4° tilted data collection to capture all possible views of the cytochrome subunits.

The final cryo-EM workflow began with patch motion correction and CTF estimation in cryoSPARC^65–67^. Subsequently, automated particle segment picking was performed using the ’Filament Tracer’ function with a 30-pixel shift between adjacent boxes. All auto-picked particles underwent multiple rounds of 2D classification, and particles belonging to poor 2D class averages were removed. Next, we aimed to first obtain a helical reconstruction of a single filament within the bundles. Potential helical symmetries were calculated from averaged power spectra generated from the raw particles (512-pixel box). Only one possibility was evident, given the clear meridian layer-line and another layer-line near the equator^68^. Next, 3D reconstruction was performed using “Helical Refinement,” followed by “Local CTF refinement,” and another round of “Helical Refinement” using the CTF-refined particles. The resolution of the helical reconstruction was estimated to be approximately 3.4 Å using Map:Map FSC.

The particles and volume obtained after helical refinement were then used as initial particles and references for 3D classification without any masking or applied symmetry. Additional filaments making contact with the central filament used as a reference began to emerge. Four out of ten classes exhibited at least one clear additional filament, and visual inspection of these volumes indicated that all had the same inter-filament interfaces but differed in the orientations of their central filaments. These classes were then merged for one asymmetric reconstruction, and the final resolution reached 3.1 Å, as estimated by Map:Map FSC. Maps used in the resolution estimation were sharpened using DeepEMHancer^69^, available in cryoSPARC. The relevant statistics are listed in Supplemental Table 1.

### Model building of cytochrome bundles

Model building began with identifying the correct cytochrome building block from the experimental cryo-EM map. From the helical reconstruction map, we used ModelAngelo^49^ to successfully trace most of the protein backbone, and the predicted protein sequence was used to perform a BLAST search against the *D. soudanensis* WTL proteome. An OmcE homolog, UniProt ID A0A0M4DKA8 (ORF name: DSOUD_2954, gene name: *omcE*), was identified as the best hit, having ∼50% sequence identity to the ModelAngelo-predicted sequence. To validate this, full-length modeling was performed for DSOUD_2954 and all other OmcE-like proteins within the proteome, and DSOUD_2954 was the only protein that could fully account for the cryo-EM map (Supplemental Figure 6).

For the modeling process, the AlphaFold3-predicted model of a single *D. soudanensis* WTL OmcE cytochrome subunit initially exhibited poor geometry, with a Ramachandran favored percentage below 60%. To address this, we employed RosettaCM^70^ rebuilding scripts with heme ligands and generated over 2,000 trajectory models. A chimera model for subsequent refinement was constructed from these models, selecting regions with top geometry statistics and manually choosing those that best matched the cryo-EM map based on visual inspection. Furthermore, accurately placing heme molecules into the cryo-EM map at this resolution without prior knowledge is challenging, as all software lacks geometry restraints for amino acid to heme interactions. Therefore, to improve the refinement of heme-interacting regions at this resolution, bond/angle restraints for the heme molecule itself, His-Fe bonds, and Cys-heme thioether bonds were implemented based on geometries observed in high-resolution crystal structures such as NrfB^71^ (PDB ID: 2P0B) and NrfHA^72^ (PDB ID: 2J7A). Similar refinement strategies have been employed for previous cytochrome nanowire studies^17,18,25^. Individual cytochrome subunits were manually adjusted against the asymmetric reconstruction map in Coot^73^, and the resulting bundle model was then real-space refined^74^ against the asymmetric map. Model quality was assessed using MolProbity^75^. Refinement statistics for the WTL bundles are presented in Supplementary Table 1.

### Structural Analysis of Heme Molecules in Heme-Containing PDB Structures

All experimental structural coordinates with heme ligands were downloaded from the Protein Data Bank (PDB). Structural predictions with heme molecules were generated using the AlphaFold3^41^ online server (https://alphafoldserver.com/). All possible heme pairs were then filtered, retaining only those with a minimum distance less than or equal to 8 Å between the two porphyrin rings, using the “contact” command in UCSF ChimeraX^76^. For each qualifying pair, the rotation matrix between the two hemes was generated in ChimeraX using the “align” command. The rotation angle, θ, was either calculated from the rotation matrix or directly obtained from the command reply log within ChimeraX.

### Data Availability

The three-dimensional reconstruction generated in this study have been deposited in the Electron Microscopy Data Bank under accession codes EMD-73506. The atomic model generated in this study have been deposited in the Protein Data Bank under accession code 9YUQ.

EMD-73506 [https://www.ebi.ac.uk/pdbe/entry/emdb/EMD-73506] 9YUQ [http://doi.org/10.2210/pdb9YUQ/pdb]

## Acknowledgements

This research was, in part, supported by the National Cancer Institute’s National Cryo-EM Facility at the Frederick National Laboratory for Cancer Research under contract 75N91019D00024. Electron microscopy screening was carried out in the UAB Cryo-EM Facility, supported by the Institutional Research Core Program and O’Neal Comprehensive Cancer Center (NIH grant P30 CA013148), with additional funding from NIH grant S10 OD024978. We are grateful to Dr. James Kizziah and Dr. Tara Fox for assisting with the screening or data collection. The work in was supported by DoE grants SC0024303 (F.W.), SC0020322 (A.I.H), SC0020329 (D.R.B.)

## Author Contributions

C.H.C. and H.A.P. performed extracellular appendage preparation and enrichment. H.A.P., S.T.R., A.Z., J.L.F. and F.W. performed microscopy image analysis. H.A.P., S.T.R., A.Z. and F.W. performed structure-based analysis. D.R.B. performed the gene cluster analysis. G.O.C., M.Z.T., performed AFM measurements. A.I.H., D.R.B. and F.W. obtained funding and supervised the research. H.A.P., A.I.H., D.R.B. and F.W. wrote the manuscript with input from all authors.

---

**Supplementary Figure 1.**
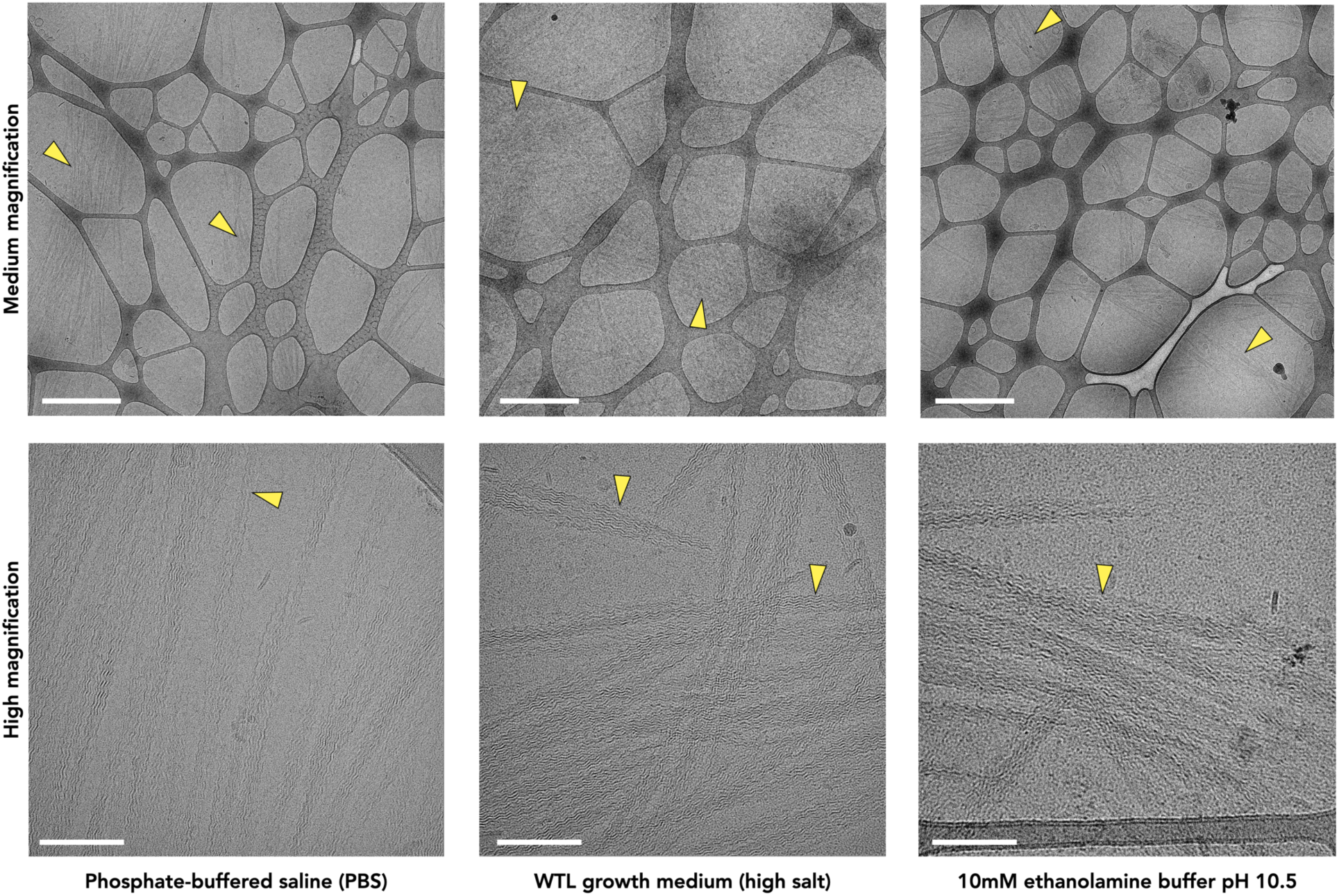
Sheared D. soudanensis nanowires in different buffers. To ensure that the formed cytochrome nanowire bundles are not an artifact of buffer conditions, we examined the bundles in three different buffer conditions: phosphate-buffered saline (left), bacterial high-salt growth medium (middle), and 10 mM ethanolamine buffer, pH 10.5 (right). The scale bar in the medium magnification is 1 µm, and the scale bars in the high magnification are 100 nm. The bundles are indicated with yellow arrowheads.

**Supplementary Figure 2.**
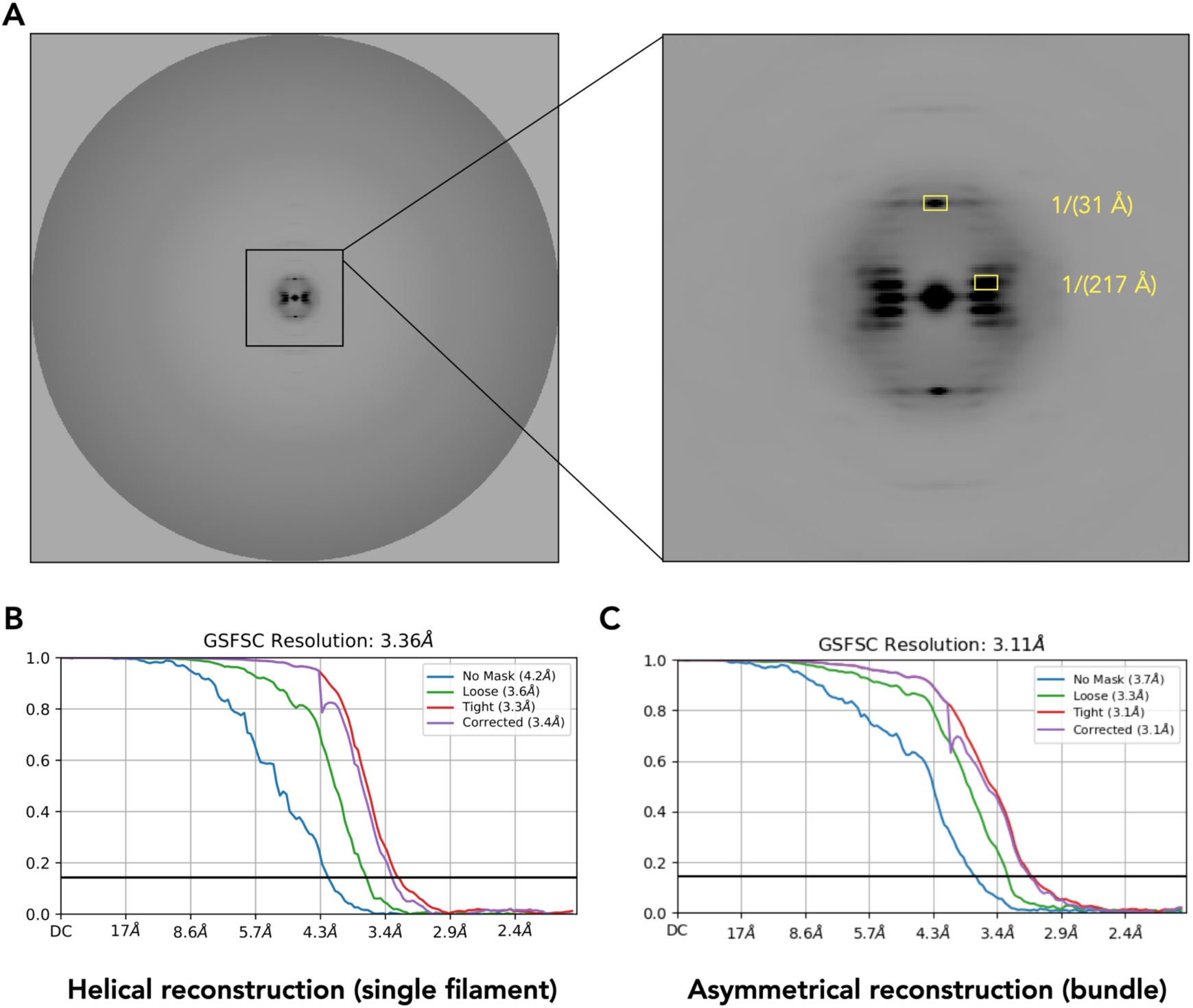
Average Power Spectrum and Fourier Shell Correlation (FSC) Calculations. **A.** The average power spectrum of nanowire raw particles aligned to the same axis. **B.** The map-to-map FSC calculation of the helical reconstruction of *D. soudanensis* cytochrome nanowire (0.143 cutoff). **C.** The map-to-map FSC calculation of the asymmetrical reconstruction of *D. soudanensis* cytochrome nanowire bundles (0.143 cutoff).

**Supplementary Figure 3.**
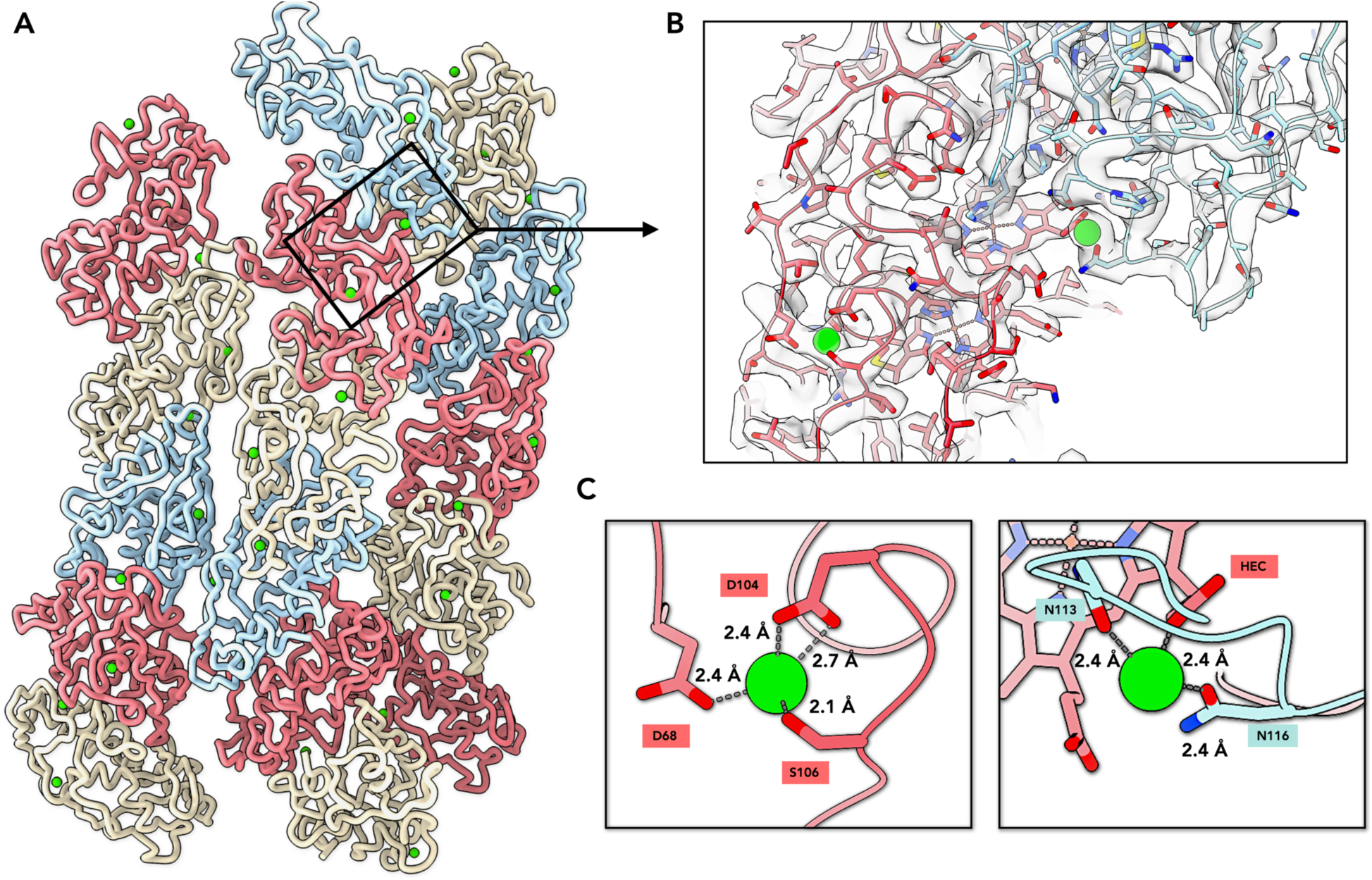
Ion Binding Sites in Cytochrome Nanowire Bundles. **A.** Ribbon view of the cytochrome nanowire bundles, with adjacent protein subunits colored differently. Putative calcium ions are displayed as green spheres. **B.** A zoomed-in view showing the putative calcium ions within the cryo-EM map. Protein backbones are shown as ribbons, while side chains are displayed as sticks. **C.** Contacts between the putative calcium ions and nearby oxygen atoms (O-atoms) from adjacent residues or a heme c molecule. Water molecules are typically invisible at this resolution.

**Supplementary Figure 4.**
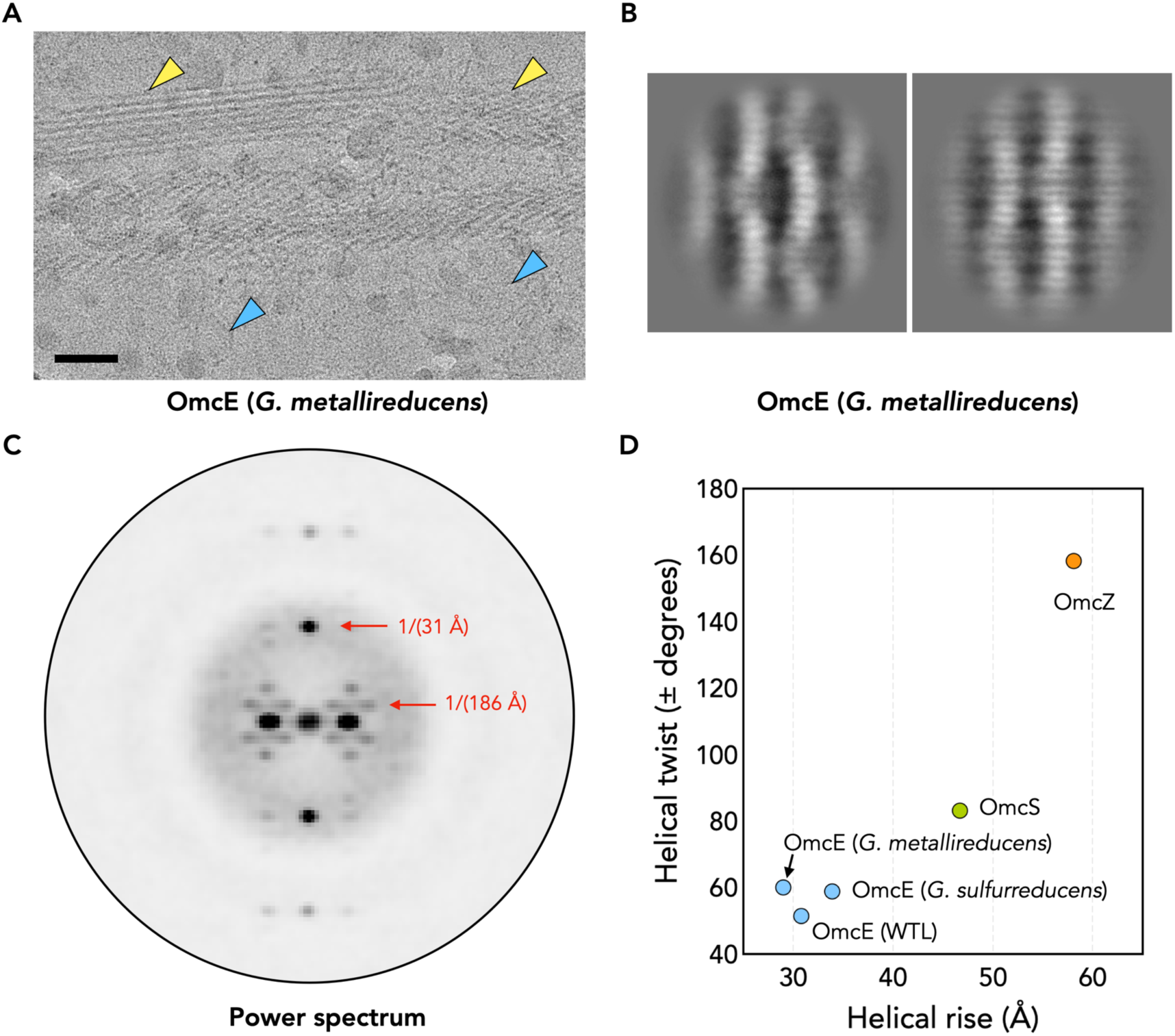
Cryo-EM of Putative OmcE Bundles in *Geobacter metallireducens*. **A.** Cryo-EM micrograph of cytochrome bundles from *Geobacter metallireducens*. Bundles are indicated by yellow arrowheads. Single fibers disassociated from the bundle are indicated with blue arrowheads. Scale bar: 50 nm. **B.** 2D averages of cytochrome bundles from *Geobacter metallireducens*. **C.** Indexing of the average power spectrum from aligned *Geobacter metallireducens* cytochrome bundles. **D.** The helical symmetry of the bundles in *Geobacter metallireducens*, compared to other bacterial cytochrome filaments, including other OmcE proteins, OmcS, and OmcZ.

**Supplementary Figure 5.**
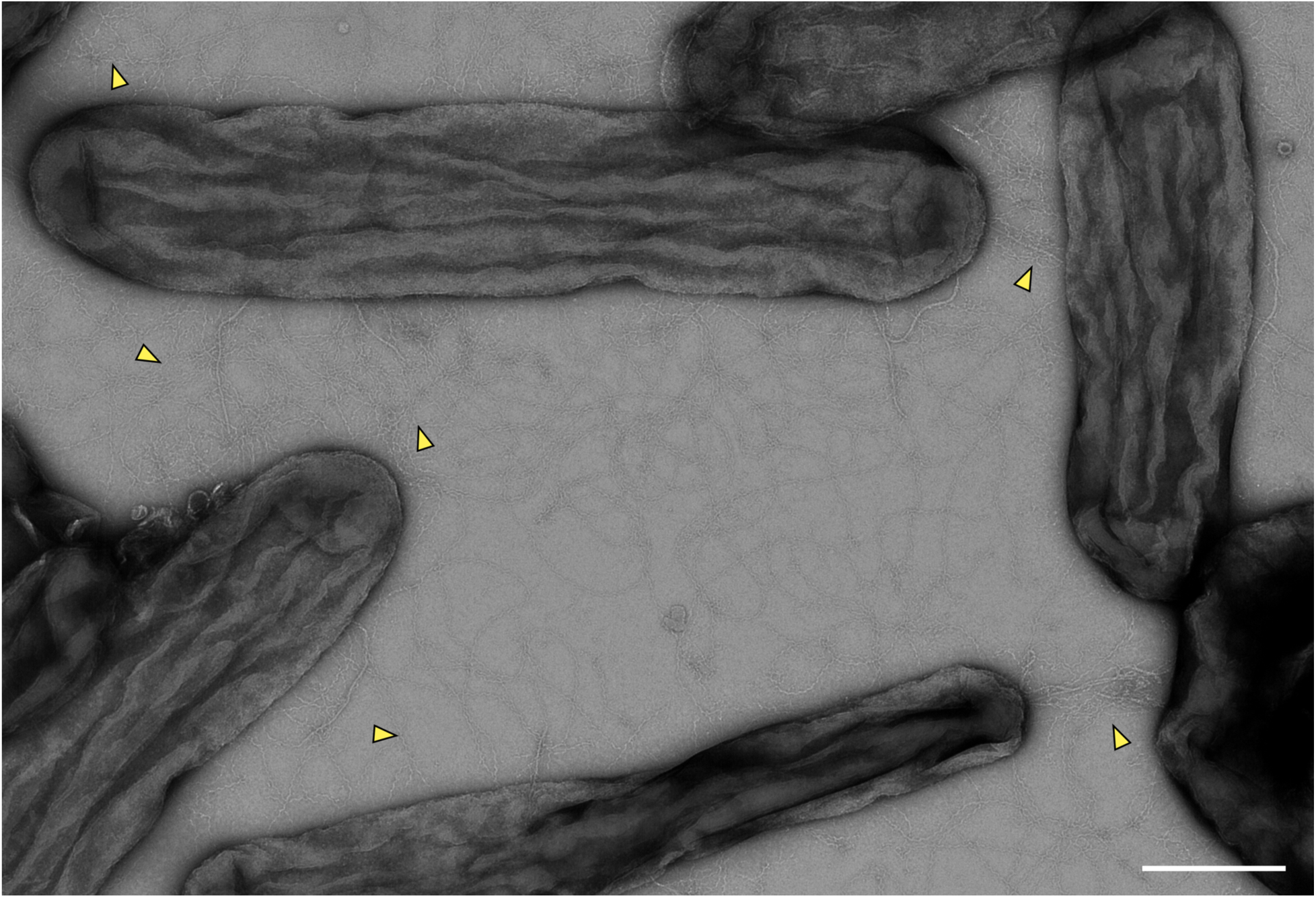
Negative staining of *D. soudanensis* cells. Negative staining images of *D. soudanensis* WTL cells grown in high salt medium. To obtain a clear view of the filaments, cells from the cultured medium, instead of biofilm aggregates, was applied to the grid. Cytochrome bundles remained visible between cells, as indicated by yellow arrowheads. Scale bar: 400 nm.

**Supplementary Figure 6.**
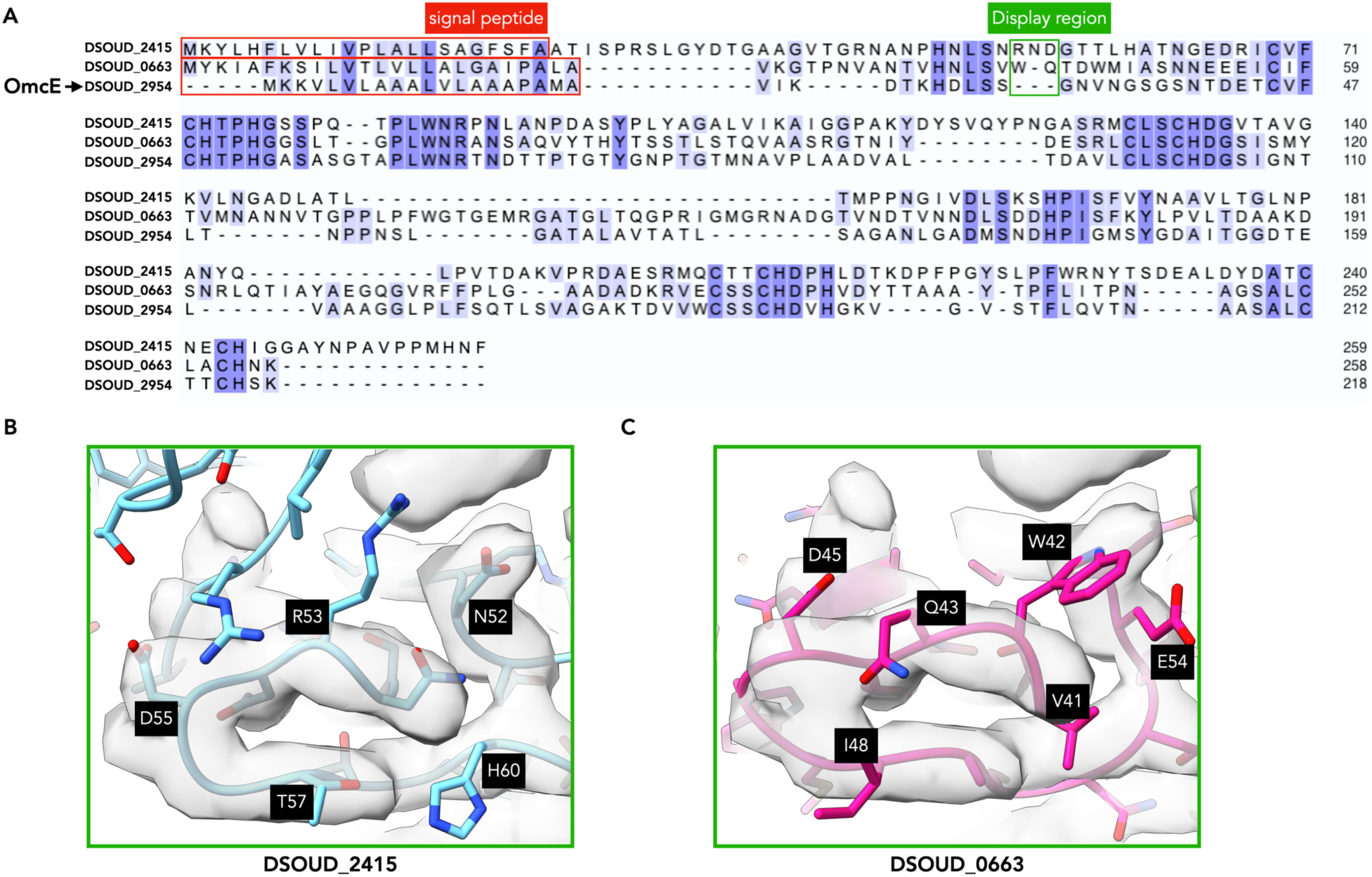
*De novo* atomic model building of OmcE filaments in *D. soudanensis* WTL. **A.** Sequence-based alignment of OmcE (DSOUD_2954) and two other c-type cytochromes (DSOUD_2415 and DSOUD_0663) with a similar protein fold. **B.** and **C**. The zoomed views of one example region indicated in (**A**) by a green box, showing the atomic model of the other two OmcE-like proteins fit into the cryo-EM map. Disagreements between the map and incorrect sequences are obvious.

**Supplementary Table 1.** Cryo-EM and refinement statistics of *Desulfuromonas soudanensis* WTL OmcE-like cytochrome nanowire.

| Parameter |  |
| --- | --- |
| **Data collection and processing** |  |
| Voltage (kV) | 300 |
| Electron exposure (e<sup>-</sup> Å<sup>-2</sup>) | 50 |
| Pixel size (Å) | 1.07 |
| Particle images (n) | 1,241,521 |
| Shift (pixel) | 30 |
| **Helical symmetry**, single filament |  |
| Point group | C1 |
| Helical rise (Å) | 30.76 |
| Helical twist (°) | 51.44 |
| **Map resolution (Å)**, asymmetrical reconstruction |  |
| Map:map FSC (0.143) | 3.1 |
| Model:map FSC (0.5) | 3.4 |
| **Refinement and Model validation** |  |
| Ramachandran Favored (%) | 88.7 |
| Ramachandran Outliers (%) | 0.4 |
| RSCC | 0.76 |
| Clashscore | 22.2 |
| Bonds RMSD, length (Å) | 0.006 |
| Bonds RMSD, angles (°) | 1.399 |
| **Deposition ID** |  |
| PDB (model) | 9YUQ |
| EMDB (map) | EMD-73506 |

## REFERENCE

1. Lovley, D.R., and Holmes, D.E. (2022). Electromicrobiology: the ecophysiology of phylogenetically diverse electroactive microorganisms. Nat Rev Microbiol 20, 5–19. 10.1038/s41579-021-00597-6.

2. Shi, L., Dong, H.L., Reguera, G., Beyenal, H., Lu, A.H., Liu, J., Yu, H.Q., and Fredrickson, J.K. (2016). Extracellular electron transfer mechanisms between microorganisms and minerals. Nature Reviews Microbiology 14, 651–662. 10.1038/nrmicro.2016.93.

3. Myers, C.R., and Nealson, K.H. (1988). Bacterial Manganese Reduction and Growth with Manganese Oxide as the Sole Electron Acceptor. Science 240, 1319–1321. doi:10.1126/science.240.4857.1319.

4. Lovley, D.R., and Phillips, E.J. (1988). Novel mode of microbial energy metabolism: organic carbon oxidation coupled to dissimilatory reduction of iron or manganese. Appl Environ Microbiol 54, 1472–1480. 10.1128/aem.54.6.1472-1480.1988.

5. Summers, Z.M., Fogarty, H.E., Leang, C., Franks, A.E., Malvankar, N.S., and Lovley, D.R. (2010). Direct Exchange of Electrons Within Aggregates of an Evolved Syntrophic Coculture of Anaerobic Bacteria. Science 330, 1413–1415. doi:10.1126/science.1196526.

6. Reguera, G. (2018). Microbial nanowires and electroactive biofilms. FEMS Microbiol Ecol 94. 10.1093/femsec/fiy086.

7. Edwards, M.J., Richardson, D.J., Paquete, C.M., and Clarke, T.A. (2020). Role of multiheme cytochromes involved in extracellular anaerobic respiration in bacteria. Protein Science 29, 830–842. 10.1002/pro.3787.

8. Joshi, K., Chan, C.H., and Bond, D.R. (2021). Geobacter sulfurreducens inner membrane cytochrome CbcBA controls electron transfer and growth yield near the energetic limit of respiration. Molecular Microbiology 116, 1124–1139. 10.1111/mmi.14801.

9. Ouboter, H.T., Mesman, R., Sleutels, T., Postma, J., Wissink, M., Jetten, M.S.M., Ter Heijne, A., Berben, T., and Welte, C.U. (2024). Mechanisms of extracellular electron transfer in anaerobic methanotrophic archaea. Nat. Commun. 15. 10.1038/s41467-024-45758-2.

10. Pirbadian, S., Barchinger, S.E., Leung, K.M., Byun, H.S., Jangir, Y., Bouhenni, R.A., Reed, S.B., Romine, M.F., Saffarini, D.A., Shi, L., et al. (2014). Shewanella oneidensis MR-1 nanowires are outer membrane and periplasmic extensions of the extracellular electron transport components. Proc Natl Acad Sci U S A 111, 12883–12888. 10.1073/pnas.1410551111.

11. Marsili, E., Baron, D.B., Shikhare, I.D., Coursolle, D., Gralnick, J.A., and Bond, D.R. (2008). Shewanella Secretes flavins that mediate extracellular electron transfer. Proceedings of the National Academy of Sciences of the United States of America 105, 3968–3973. 10.1073/pnas.0710525105.

12. Subramanian, P., Pirbadian, S., El-Naggar, M.Y., and Jensen, G.J. (2018). Ultrastructure of MR-1 nanowires revealed by electron cryotomography. Proceedings of the National Academy of Sciences of the United States of America 115, E3246–E3255. 10.1073/pnas.1718810115.

13. Liu, Y., Wang, Z., Liu, J., Levar, C., Edwards, M.J., Babauta, J.T., Kennedy, D.W., Shi, Z., Beyenal, H., Bond, D.R., et al. (2014). A trans-outer membrane porin-cytochrome protein complex for extracellular electron transfer by eobacter sulfurreducens PCA. Environmental Microbiology Reports 6, 776–785. 10.1111/1758-2229.12204.

14. Hartshorne, R.S., Reardon, C.L., Ross, D., Nuester, J., Clarke, T.A., Gates, A.J., Mills, P.C., Fredrickson, J.K., Zachara, J.M., Shi, L., et al. (2009). Characterization of an electron conduit between bacteria and the extracellular environment. Proc Natl Acad Sci USA 106, 22169–22174. doi:10.1073/pnas.0900086106.

15. Babauta, J., Renslow, R., Lewandowski, Z., and Beyenal, H. (2012). Electrochemically active biofilms: facts and fiction. A review. Biofouling 28, 789–812. 10.1080/08927014.2012.710324.

16. Wang, F., Craig, L., Liu, X., Rensing, C., and Egelman, E.H. (2023). Microbial nanowires: type IV pili or cytochrome filaments? Trends Microbiol 31, 384–392. 10.1016/j.tim.2022.11.004.

17. Wang, F., Mustafa, K., Suciu, V., Joshi, K., Chan, C.H., Choi, S., Su, Z., Si, D., Hochbaum, A.I., Egelman, E.H., and Bond, D.R. (2022). Cryo-EM structure of an extracellular Geobacter OmcE cytochrome filament reveals tetrahaem packing. Nat Microbiol 7, 1291–1300. 10.1038/s41564-022-01159-z.

18. Wang, F., Chan, C.H., Suciu, V., Mustafa, K., Ammend, M., Si, D., Hochbaum, A.I., Egelman, E.H., and Bond, D.R. (2022). Structure of Geobacter OmcZ filaments suggests extracellular cytochrome polymers evolved independently multiple times. Elife 11. 10.7554/eLife.81551.

19. Wang, F., Gu, Y., O’Brien, J.P., Yi, S.M., Yalcin, S.E., Srikanth, V., Shen, C., Vu, D., Ing, N.L., Hochbaum, A.I., et al. (2019). Structure of Microbial Nanowires Reveals Stacked Hemes that Transport Electrons over Micrometers. Cell 177, 361–369 e310. 10.1016/j.cell.2019.03.029.

20. Gu, Y., Guberman-Pfeffer, M.J., Srikanth, V., Shen, C., Giska, F., Gupta, K., Londer, Y., Samatey, F.A., Batista, V.S., and Malvankar, N.S. (2023). Structure of Geobacter cytochrome OmcZ identifies mechanism of nanowire assembly and conductivity. Nat Microbiol 8, 284–298. 10.1038/s41564-022-01315-5.

21. Filman, D.J., Marino, S.F., Ward, J.E., Yang, L., Mester, Z., Bullitt, E., Lovley, D.R., and Strauss, M. (2019). Cryo-EM reveals the structural basis of long-range electron transport in a cytochrome-based bacterial nanowire. Commun Biol 2, 219. 10.1038/s42003-019-0448-9.

22. Coppi, M.V., Leang, C., Sandler, S.J., and Lovley, D.R. (2001). Development of a genetic system for *Geobacter sulfurreducens*. Applied and Environmental Microbiology 67, 3180–3187. Doi 10.1128/Aem.67.7.3180-3187.2001.

23. Chan, C.H., Levar, C.E., Jimenez-Otero, F., and Bond, D.R. (2017). Genome Scale Mutational Analysis of Geobacter sulfurreducens Reveals Distinct Molecular Mechanisms for Respiration and Sensing of Poised Electrodes versus Fe(III) Oxides. J Bacteriol 199. 10.1128/JB.00340-17.

24. Chan, C.H., Levar, C.E., Zacharoff, L., Badalamenti, J.P., and Bond, D.R. (2015). Scarless Genome Editing and Stable Inducible Expression Vectors for Geobacter sulfurreducens. Appl Environ Microbiol 81, 7178–7186. 10.1128/AEM.01967-15.

25. Baquero, D.P., Cvirkaite-Krupovic, V., Hu, S.S., Fields, J.L., Liu, X., Rensing, C., Egelman, E.H., Krupovic, M., and Wang, F. (2023). Extracellular cytochrome nanowires appear to be ubiquitous in prokaryotes. Cell 186, 2853–2864 e2858. 10.1016/j.cell.2023.05.012.

26. Franks, A.E., Glaven, R.H., and Lovley, D.R. (2012). Real-Time Spatial Gene Expression Analysis within Current-Producing Biofilms. Chemsuschem 5, 1092–1098. 10.1002/cssc.201100714.

27. Nevin, K.P., Kim, B.C., Glaven, R.H., Johnson, J.P., Woodard, T.L., Methé, B.A., DiDonato, R.J., Covalla, S.F., Franks, A.E., Liu, A., and Lovley, D.R. (2009). Anode Biofilm Transcriptomics Reveals Outer Surface Components Essential for High Density Current Production in Fuel Cells. Plos One 4. 10.1371/journal.pone.0005628.

28. Peng, L., and Zhang, Y. (2017). Cytochrome OmcZ is essential for the current generation by under low electrode potential. Electrochim Acta 228, 447–452. 10.1016/j.electacta.2017.01.091.

29. Richter, H., Nevin, K.P., Jia, H.F., Lowy, D.A., Lovley, D.R., and Tender, L.M. (2009). Cyclic voltammetry of biofilms of wild type and mutant on fuel cell anodes indicates possible roles of OmcB, OmcZ, type IV pili, and protons in extracellular electron transfer. Energy & Environmental Science 2, 506–516. 10.1039/b816647a.

30. Hall-Stoodley, L., Costerton, J.W., and Stoodley, P. (2004). Bacterial biofilms: From the natural environment to infectious diseases. Nature Reviews Microbiology 2, 95–108. 10.1038/nrmicro821.

31. Vidakovic, L., Mikhaleva, S., Jeckel, H., Nisnevich, V., Strenger, K., Neuhaus, K., Raveendran, K., Ben-Moshe, N.B., Aznaourova, M., Nosho, K., et al. (2023). Biofilm formation on human immune cells is a multicellular predation strategy of Vibrio cholerae. Cell 186, 2690-+. 10.1016/j.cell.2023.05.008.

32. Drame, I., Lafforgue, C., Formosa-Dague, C., Chapot-Chartier, M.P., Piard, J.C., Castelain, M., and Dague, E. (2021). Pili and other surface proteins influence the structure and the nanomechanical properties of Lactococcus lactis biofilms. Sci Rep 11, 4846. 10.1038/s41598-021-84030-1.

33. Lemon, K.P., Higgins, D.E., and Kolter, R. (2007). Flagellar motility is critical for biofilm formation. Journal of Bacteriology 189, 4418–4424. 10.1128/Jb.01967-06.

34. Ligthart, K., Belzer, C., de Vos, W.M., and Tytgat, H.L.P. (2020). Bridging Bacteria and the Gut: Functional Aspects of Type IV Pili. Trends Microbiol 28, 340–348. 10.1016/j.tim.2020.02.003.

35. Wang, F., Cvirkaite-Krupovic, V., Krupovic, M., and Egelman, E.H. (2022). Archaeal bundling pili of Pyrobaculum calidifontis reveal similarities between archaeal and bacterial biofilms. Proc Natl Acad Sci U S A 119, e2207037119. 10.1073/pnas.2207037119.

36. Bohning, J., Ghrayeb, M., Pedebos, C., Abbas, D.K., Khalid, S., Chai, L., and Bharat, T.A.M. (2022). Donor-strand exchange drives assembly of the TasA scaffold in Bacillus subtilis biofilms. Nat Commun 13, 7082. 10.1038/s41467-022-34700-z.

37. Bohning, J., Dobbelstein, A.W., Sulkowski, N., Eilers, K., von Kugelgen, A., Tarafder, A.K., Peak-Chew, S.Y., Skehel, M., Alva, V., Filloux, A., and Bharat, T.A.M. (2023). Architecture of the biofilm-associated archaic Chaperone-Usher pilus CupE from Pseudomonas aeruginosa. PLoS Pathog 19, e1011177. 10.1371/journal.ppat.1011177.

38. Doucey, M.A., and Carrara, S. (2019). Nanowire Sensors in Cancer. Trends Biotechnol 37, 86–99. 10.1016/j.tibtech.2018.07.014.

39. Wang, Z., Lee, S., Koo, K., and Kim, K. (2016). Nanowire-Based Sensors for Biological and Medical Applications. IEEE Trans Nanobioscience 15, 186–199. 10.1109/TNB.2016.2528258.

40. Domigan, L.J. (2013). Proteins and Peptides as Biological Nanowires: Towards Biosensing Devices. In Protein Nanotechnology: Protocols, Instrumentation, and Applications, Second Edition, J.A. Gerrard, ed. (Humana Press), pp. 131–152. 10.1007/978-1-62703-354-1_8.

41. Abramson, J., Adler, J., Dunger, J., Evans, R., Green, T., Pritzel, A., Ronneberger, O., Willmore, L., Ballard, A.J., Bambrick, J., et al. (2024). Accurate structure prediction of biomolecular interactions with AlphaFold 3. Nature. 10.1038/s41586-024-07487-w.

42. Badalamenti, J.P., Summers, Z.M., Chan, C.H., Gralnick, J.A., and Bond, D.R. (2016). Isolation and Genomic Characterization of ‘Desulfuromonas soudanensis WTL’, a Metal- and Electrode-Respiring Bacterium from Anoxic Deep Subsurface Brine. Frontiers in Microbiology 7. 10.3389/fmicb.2016.00913.

43. Wang, F., Coureuil, M., Osinski, T., Orlova, A., Altindal, T., Gesbert, G., Nassif, X., Egelman, E.H., and Craig, L. (2017). Cryoelectron Microscopy Reconstructions of the Pseudomonas aeruginosa and Neisseria gonorrhoeae Type IV Pili at Sub-nanometer Resolution. Structure 25, 1423–1435 e1424. 10.1016/j.str.2017.07.016.

44. Bazayeva, M., Andreini, C., and Rosato, A. (2024). A database overview of metal-coordination distances in metalloproteins. Acta Crystallogr D Struct Biol 80, 362–376. 10.1107/S2059798324003152.

45. Krissinel, E., and Henrick, K. (2007). Inference of macromolecular assemblies from crystalline state. J Mol Biol 372, 774–797. 10.1016/j.jmb.2007.05.022.

46. Childers, S.E., Ciufo, S., and Lovley, D.R. (2002). Geobacter metallireducens accesses insoluble Fe(III) oxide by chemotaxis. Nature 416, 767–769. 10.1038/416767a.

47. Rotaru, A.E., Shrestha, P.M., Liu, F., Markovaite, B., Chen, S., Nevin, K.P., and Lovley, D.R. (2014). Direct Interspecies Electron Transfer between Geobacter metallireducens and Methanosarcina barkeri Applied and Environmental Microbiology 80, 4599–4605. 10.1128/Aem.00895-14.

48. Kuhlbrandt, W. (2014). Biochemistry. The resolution revolution. Science 343, 1443–1444. 10.1126/science.1251652.

49. Jamali, K., Kall, L., Zhang, R., Brown, A., Kimanius, D., and Scheres, S.H.W. (2024). Automated model building and protein identification in cryo-EM maps. Nature 628, 450–457. 10.1038/s41586-024-07215-4.

50. Pfab, J., Phan, N.M., and Si, D. (2021). DeepTracer for fast de novo cryo-EM protein structure modeling and special studies on CoV-related complexes. Proc Natl Acad Sci U S A 118. 10.1073/pnas.2017525118.

51. Wang, T., Li, Z., Xu, K., Huang, W., Huang, G., Zhang, Q.C., and Yan, N. (2024). CryoSeek: A strategy for bioentity discovery using cryoelectron microscopy. Proc Natl Acad Sci U S A 121, e2417046121. 10.1073/pnas.2417046121.

52. Semchonok, D.A., Kyrilis, F.L., Hamdi, F., and Kastritis, P.L. (2023). Cryo-EM of a heterogeneous biochemical fraction elucidates multiple protein complexes from a multicellular thermophilic eukaryote. J Struct Biol X 8, 100094. 10.1016/j.yjsbx.2023.100094.

53. Liu, J., Eastep, G.N., Cvirkaite-Krupovic, V., Rich-New, S.T., Kreutzberger, M.A.B., Egelman, E.H., Krupovic, M., and Wang, F. (2024). Two distinct archaeal type IV pili structures formed by proteins with identical sequence. Nat Commun 15, 5049. 10.1038/s41467-024-45062-z.

54. Voica, D.M., Bartha, L., Banciu, H.L., and Oren, A. (2016). Heavy metal resistance in halophilic Bacteria and Archaea. FEMS Microbiol Lett 363. 10.1093/femsle/fnw146.

55. Nies, D.H. (2003). Efflux-mediated heavy metal resistance in prokaryotes. FEMS Microbiol Rev 27, 313–339. 10.1016/S0168-6445(03)00048-2.

56. Liu, J., Chakraborty, S., Hosseinzadeh, P., Yu, Y., Tian, S.L., Petrik, I., Bhagi, A., and Lu, Y. (2014). Metalloproteins Containing Cytochrome, Iron-Sulfur, or Copper Redox Centers. Chemical Reviews 114, 4366–4469. 10.1021/cr400479b.

57. Pakharukova, N., Malmi, H., Tuittila, M., Dahlberg, T., Ghosal, D., Chang, Y.W., Myint, S.L., Paavilainen, S., Knight, S.D., Lamminmäki, U., et al. (2022). Archaic chaperone-usher pili self-secrete into superelastic zigzag springs. Nature 609, 335-+. 10.1038/s41586-022-05095-0.

58. Nuccio, S.P., and Bäumler, A.J. (2007). Evolution of the chaperone/usher assembly pathway:: Fimbrial classification goes Greek. Microbiol Mol Biol R 71, 551-+. 10.1128/Mmbr.00014-07.

59. Vallet, I., Olson, J.W., Lory, S., Lazdunski, A., and Filloux, A. (2001). The chaperone/usher pathways of *Pseudomonas aeruginosa*: Identification of fimbrial gene clusters (cup) and their involvement in biofilm formation. Proceedings of the National Academy of Sciences of the United States of America 98, 6911–6916. DOI 10.1073/pnas.111551898.

60. Gu, Y., Srikanth, V., Salazar-Morales, A.I., Jain, R., O’Brien, J.P., Yi, S.M., Soni, R.K., Samatey, F.A., Yalcin, S.E., and Malvankar, N.S. (2021). Structure of Geobacter pili reveals secretory rather than nanowire behaviour. Nature 597, 430–434. 10.1038/s41586-021-03857-w.

61. Shen, C., Salazar-Morales, A.I., Jung, W., Erwin, J., Gu, Y., Coelho, A., Gupta, K., Yalcin, S.E., Samatey, F.A., and Malvankar, N.S. (2025). A widespread and ancient bacterial machinery assembles cytochrome OmcS nanowires essential for extracellular electron transfer. Cell Chem Biol 32, 239–254 e237. 10.1016/j.chembiol.2024.12.013.

62. Allen, W.J., Phan, G., and Waksman, G. (2012). Pilus biogenesis at the outer membrane of Gram-negative bacterial pathogens. Curr Opin Struc Biol 22, 500–506. 10.1016/j.sbi.2012.02.001.

63. Werneburg, G.T., and Thanassi, D.G. (2018). Pili Assembled by the Chaperone/Usher Pathway in Escherichia coli and Salmonella. EcoSal Plus 8. 10.1128/ecosalplus.ESP-0007-2017.

64. Busch, A., and Waksman, G. (2012). Chaperone-usher pathways: diversity and pilus assembly mechanism. Philos T R Soc B 367, 1112–1122. 10.1098/rstb.2011.0206.

65. Zheng, S.Q., Palovcak, E., Armache, J.P., Verba, K.A., Cheng, Y., and Agard, D.A. (2017). MotionCor2: anisotropic correction of beam-induced motion for improved cryo-electron microscopy. Nat Methods 14, 331–332. 10.1038/nmeth.4193.

66. Punjani, A., Rubinstein, J.L., Fleet, D.J., and Brubaker, M.A. (2017). cryoSPARC: algorithms for rapid unsupervised cryo-EM structure determination. Nat Methods 14, 290–296. 10.1038/nmeth.4169.

67. Rohou, A., and Grigorieff, N. (2015). CTFFIND4: Fast and accurate defocus estimation from electron micrographs. J Struct Biol 192, 216–221. 10.1016/j.jsb.2015.08.008.

68. Wang, F., Gnewou, O., Solemanifar, A., Conticello, V.P., and Egelman, E.H. (2022). Cryo-EM of Helical Polymers. Chem Rev 122, 14055–14065. 10.1021/acs.chemrev.1c00753.

69. Sanchez-Garcia, R., Gomez-Blanco, J., Cuervo, A., Carazo, J.M., Sorzano, C.O.S., and Vargas, J. (2021). DeepEMhancer: a deep learning solution for cryo-EM volume post-processing. Commun Biol 4, 874. 10.1038/s42003-021-02399-1.

70. Wang, R.Y., Kudryashev, M., Li, X., Egelman, E.H., Basler, M., Cheng, Y., Baker, D., and DiMaio, F. (2015). De novo protein structure determination from near-atomic-resolution cryo-EM maps. Nat Methods 12, 335–338. 10.1038/nmeth.3287.

71. Clarke, T.A., Cole, J.A., Richardson, D.J., and Hemmings, A.M. (2007). The crystal structure of the pentahaem-type cytochrome NrfB and characterization of its solution-state interaction with the pentahaem nitrite reductase NrfA. Biochemical Journal 406, 19–30. 10.1042/Bj20070321.

72. Rodrigues, M.L., Oliveira, T.F., Pereira, I.A.C., and Archer, M. (2006). X-ray structure of the membrane-bound cytochrome quinol dehydrogenase NrfH reveals novel haem coordination. Embo Journal 25, 5951–5960. 10.1038/sj.emboj.7601439.

73. Emsley, P., and Cowtan, K. (2004). Coot: model-building tools for molecular graphics. Acta Crystallogr D Biol Crystallogr 60, 2126–2132. 10.1107/S0907444904019158.

74. Afonine, P.V., Poon, B.K., Read, R.J., Sobolev, O.V., Terwilliger, T.C., Urzhumtsev, A., and Adams, P.D. (2018). Real-space refinement in PHENIX for cryo-EM and crystallography. Acta Crystallogr D Struct Biol 74, 531–544. 10.1107/S2059798318006551.

75. Williams, C.J., Headd, J.J., Moriarty, N.W., Prisant, M.G., Videau, L.L., Deis, L.N., Verma, V., Keedy, D.A., Hintze, B.J., Chen, V.B., et al. (2018). MolProbity: More and better reference data for improved all-atom structure validation. Protein Sci 27, 293–315. 10.1002/pro.3330.

76. Pettersen, E.F., Goddard, T.D., Huang, C.C., Meng, E.C., Couch, G.S., Croll, T.I., Morris, J.H., and Ferrin, T.E. (2021). UCSF ChimeraX: Structure visualization for researchers, educators, and developers. Protein Sci 30, 70–82. 10.1002/pro.3943.

